# Broadly neutralizing antibody induction by non-stabilized SARS-CoV-2 Spike mRNA vaccination in nonhuman primates

**DOI:** 10.1101/2023.12.18.572191

**Authors:** R. Dilshan Malewana, Victoria Stalls, Aaron May, Xiaozhi Lu, David R. Martinez, Alexandra Schäfer, Dapeng Li, Maggie Barr, Laura L. Sutherland, Esther Lee, Robert Parks, Whitney Edwards Beck, Amanda Newman, Kevin W. Bock, Mahnaz Minai, Bianca M. Nagata, C. Todd DeMarco, Thomas N. Denny, Thomas H. Oguin, Wes Rountree, Yunfei Wang, Katayoun Mansouri, Robert J. Edwards, Gregory D. Sempowski, Amanda Eaton, Hiromi Muramatsu, Rory Henderson, Ying Tam, Christopher Barbosa, Juanjie Tang, Derek W. Cain, Sampa Santra, Ian N. Moore, Hanne Andersen, Mark G. Lewis, Hana Golding, Robert Seder, Surender Khurana, David C. Montefiori, Norbert Pardi, Drew Weissman, Ralph S. Baric, Priyamvada Acharya, Barton F. Haynes, Kevin O. Saunders

## Abstract

Immunization with mRNA or viral vectors encoding spike with diproline substitutions (S-2P) has provided protective immunity against severe COVID-19 disease. How immunization with Severe Acute Respiratory Syndrome Coronavirus 2 (SARS-CoV-2) spike elicits neutralizing antibodies (nAbs) against difficult-to-neutralize variants of concern (VOCs) remains an area of great interest. Here, we compare immunization of macaques with mRNA vaccines expressing ancestral spike either including or lacking diproline substitutions, and show the diproline substitutions were not required for protection against SARS-CoV-2 challenge or induction of broadly neutralizing B cell lineages. One group of nAbs elicited by the ancestral spike lacking diproline substitutions targeted the outer face of the receptor binding domain (RBD), neutralized all tested SARS-CoV-2 VOCs including Omicron XBB.1.5, but lacked cross-Sarbecovirus neutralization. Structural analysis showed that the macaque broad SARS-CoV-2 VOC nAbs bound to the same epitope as a human broad SARS-CoV-2 VOC nAb, DH1193. Vaccine-induced antibodies that targeted the RBD inner face neutralized multiple Sarbecoviruses, protected mice from bat CoV RsSHC014 challenge, but lacked Omicron variant neutralization. Thus, ancestral SARS-CoV-2 spike lacking proline substitutions encoded by nucleoside-modified mRNA can induce B cell lineages binding to distinct RBD sites that either broadly neutralize animal and human Sarbecoviruses or recent Omicron VOCs.

**One Sentence Summary:** Non-stabilized SARS-CoV-2 Spike mRNA vaccination activated B cells that target either conserved epitopes on SARS-CoV-2 Omicron variants of concern, or cross-neutralizing epitopes on pre-emergent Sarbecoviruses.

## INTRODUCTION

As of November 2023, the COVID-19 pandemic has claimed approximately 7 million lives globally, and the emergence of new SARS-CoV-2 variants continues to be a challenge (*1*). Neutralizing antibodies (nAbs) from active immunization remain protective immune correlates against SARS-CoV-2 infection (*2–7*). The target for nAbs against SARS-CoV-2 is the spike (S) protein on the virion surface (*8, 9*). The SARS-CoV-2 S is a trimeric type I fusion protein (*9–13*) cleaved by proteases into S1 and S2 domains. The receptor binding domain (RBD), located within the S1 domain of spike, engages the viral receptor ACE2 on target cells (14). The S2 domain mediates fusion with the target cell membrane (*11–14*). The spike trimer has a dynamic structure, where the receptor binding domain can adopt various positions (*15*). When engaging the host receptor ACE2, the RBD is in the “up” conformation. The RBD stochastically samples the “up” and “down” positions (*16, 17*). Hence, the RBD on individual protomers of the trimeric S protein can adopt different positions where all three or only a subset of RBDs can be in the up position (*9, 10*).

Over half a billion doses of mRNA encapsulated in lipid nanoparticle (mRNA-LNP) vaccines from Pfizer and Moderna have been administered in the United States (*18*). Initially, each of these vaccines encoded the S protein from the ancestral Wuhan Hu-1 or WA-1 isolates that was presumed to be stabilized in the prefusion state by including diproline substitutions K986P and V987P (S-2P) (*19*). These same substitutions improved neutralizing antibody elicitation of SARS-CoV-1 and MERS-CoV, and thus were thought to function similarly in the context of SARS-CoV-2 (*20, 21*). However, side-by-side biochemical analyses revealed the 2P mutations did not increase expression of trimeric SARS-CoV-2 Spike protein, thermostability, or antigenicity (*22*). Moreover, cryo-electron microscopy studies showed the overall structures of S with or without the diproline substitutions to be highly similar with an overall root mean square deviation of 0.54(*22*). In mice, the 2P substitutions were not required for potent immunogenicity of SARS-CoV-2 spike (*23*). It should also be noted that the Oxford-AstraZeneca COVID-19 vaccine developers chose to express from an adenoviral vector spike without the 2P substitutions for their vaccine (*24, 25*). Thus, despite the successes of the authorized mRNA-LNP vaccines encoding S-2P, the effects of the 2P substitutions on immunogenicity of SARS-CoV-2 S remains unclear.

The relationship between immunogenicity of RBD relative to its conformational dynamics has yet to be fully explored but could inform the design of next generation COVID-19 vaccines (*23*). Cryo-electron microscopy structures of S protein with or without 2P substitutions showed particles with either 1 RBD up or 3 RBDs up (*10, 17, 21, 22, 26, 27*). Previously, we designed a spike with a *de novo* disulfide bond that prevented the RBD from transitioning to the up state (*17*). Structural studies of this stabilized version of SARS-CoV-2 S showed all three RBDs were most frequently in the down position (*17*). While RBD neutralizing antibodies such as DH1041 (RBD-1 community) or the RBD inner face Ab DH1047 (RBD-6/7 community), which spans across RBD-6 and 7 epitopes, bind to the up conformation of RBD, it is unknown if an S with RBDs usually in the down state would elicit RBD neutralizing antibody responses.

Sarbecoviruses circulating in bats that have the ability to infect primary airway human cells *in vitro* are considered pre-emergent threats for zoonotic transmission (*28, 29*). *Sarbecovirus* cross-nAb DH1047 protects against bat coronavirus infection (*30, 31*), but is unable to neutralize Omicron VOCs (*32*). The escape of the Omicron sublineage from the RBD-6/7 community antibody DH1047 raises the question of whether vaccination or infection can elicit a single antibody that simultaneously neutralizes multiple zoonotic SARS-related Sarbecoviruses as well as recent SARS-CoV-2 VOCs.

Here, we show that 2P stabilization of transmembrane spike was not required for mRNA-LNP elicitation of either broad sarbecovirus or broad SARS-CoV-2 VOC nAbs. Rather both types of SARS-CoV-2 bnAbs were induced by mRNA-encoded spike lacking 2P substitutions. Such a polyclonal response provides one mechanism for how SARS-CoV-2 spike can induce broad protective antibodies against SARS-CoV-2 VOCs as well as bat zoonotic pre-emergent Sarbecoviruses.

## RESULTS

### Immunogenicity of engineered and wildtype SARS-CoV-2 spike

To investigate the effects of spike stabilization on immunogenicity and protection from infection in non-human primates, we generated nucleoside-modified mRNA-LNP vaccines encoding Wuhan-Hu-1 full-length transmembrane spike with two different potential stabilization strategies. First, the spike was stabilized with the diproline substitutions and termed S-2P. Second, the diproline substitutions were introduced into spike in combination with S483C and D985C (S-2C 2P)substitutions. Recombinant S-2C 2P has been previously shown to have RBDs predominantly in the down position (*17*). For comparison, transmembrane spike lacking any stabilization substitutions (S-tm) was also designed as an mRNA. Each of the three spike designs had the furin cleavage site replaced with a glycine-serine-alanine linker. We also generated an mRNA-LNP encoding the Wuhan-Hu-1 RBD to compare bnAb elicitation to non-stabilized (*33*) and stabilized full-length spike (**Fig. 1A**). As a negative control, we produced mRNA-LNP vaccines expressing firefly luciferase (Luc) protein instead of coronavirus S protein. Groups of eight rhesus macaques were immunized 2 times four weeks apart with 100 mcg of mRNA-LNPs encoding one of the spike proteins or the control mRNA encoding Luc (**Fig. 1B**). All rhesus macaques, except those in the control group, elicited IgG responses to Wuhan-Hu-1 S-2P, RBD, spike N-terminal domain (NTD), and S2 domain after one immunization, with titers being boosted by the second immunization (**Fig. 1C, fig. S1**). Significantly lower S-2P IgG was present for the monomeric RBD mRNA-LNP group after the first immunization (**Fig. 1C**; Exact Wilcoxon *P*<0.05 n = 8 macaques), which may indicate a low level of *in vivo* expression upon vaccination. RBD-specific serum IgG binding was highest for S-tm after one immunization (Exact Wilcoxon *P*<0.05 n = 8 macaques), but after two immunizations binding magnitudes were similar in all full-length S and monomer RBD mRNA-LNP groups (**Fig. 1C**). Similarly, after two immunizations, the serum dilution that neutralized 50% (ID_50_) of SARS-CoV-2 Wuhan-Hu1 live virus or pseudovirus replication was comparable across the different spike-immunized groups (**Fig. 1D**). We next performed competitive ELISA binding assays in which macaque plasma was used to block the binding of ACE2, RBD nAbs, and N-terminal domain nAbs. Each S-immunized group demonstrated comparable plasma blocking of ACE-2 **(Fig. 1E).** Compared to RBD mRNA-LNP immunization, S-tm mRNA-LNP elicited significantly higher antibodies that blocked ancestral SARS-CoV-2-specific RBD nAb DH1041 that target the RBD-1 community epitope, RBD inner face, *Sarbecovirus* cross-nAb DH1047 that binds across RBD-6 and 7 epitopes (Fig. 1E and **fig. S2**, Exact Wilcoxon P= P<0.05 for each antibody). SARS-CoV-2 VOCs broadly neutralizing antibody DH1193, which binds the RBD-4 community epitope (**fig. S2**), was also blocked to similar magnitudes by plasma antibodies from the various spike mRNA-LNP vaccinated macaques (**Fig. 1E**). The S-2P and S-2P 2C groups were not significantly different from each other (Exact Wilcoxon *P*>0.05 n = 8 macaques) (**Fig. 1E**). Similarly, macaque plasma from S-tm-immunized macaques blocked ancestral SARS-CoV-2 NTD nAb DH1050.1 and non-neutralizing NTD antibody DH1052 significantly better than plasma from other immunization groups (Exact Wilcoxon *P*<0.05 n = 8 macaques, **Fig. 1E**). Thus, nucleoside-modified mRNA-LNP vaccines encoding various forms of transmembrane S protein elicited similar serum neutralization responses, with S-tm generating superior antibodies that block neutralizing epitopes (Exact Wilcoxon *P*<0.05 n = 8 macaques).

**Figure 1.**
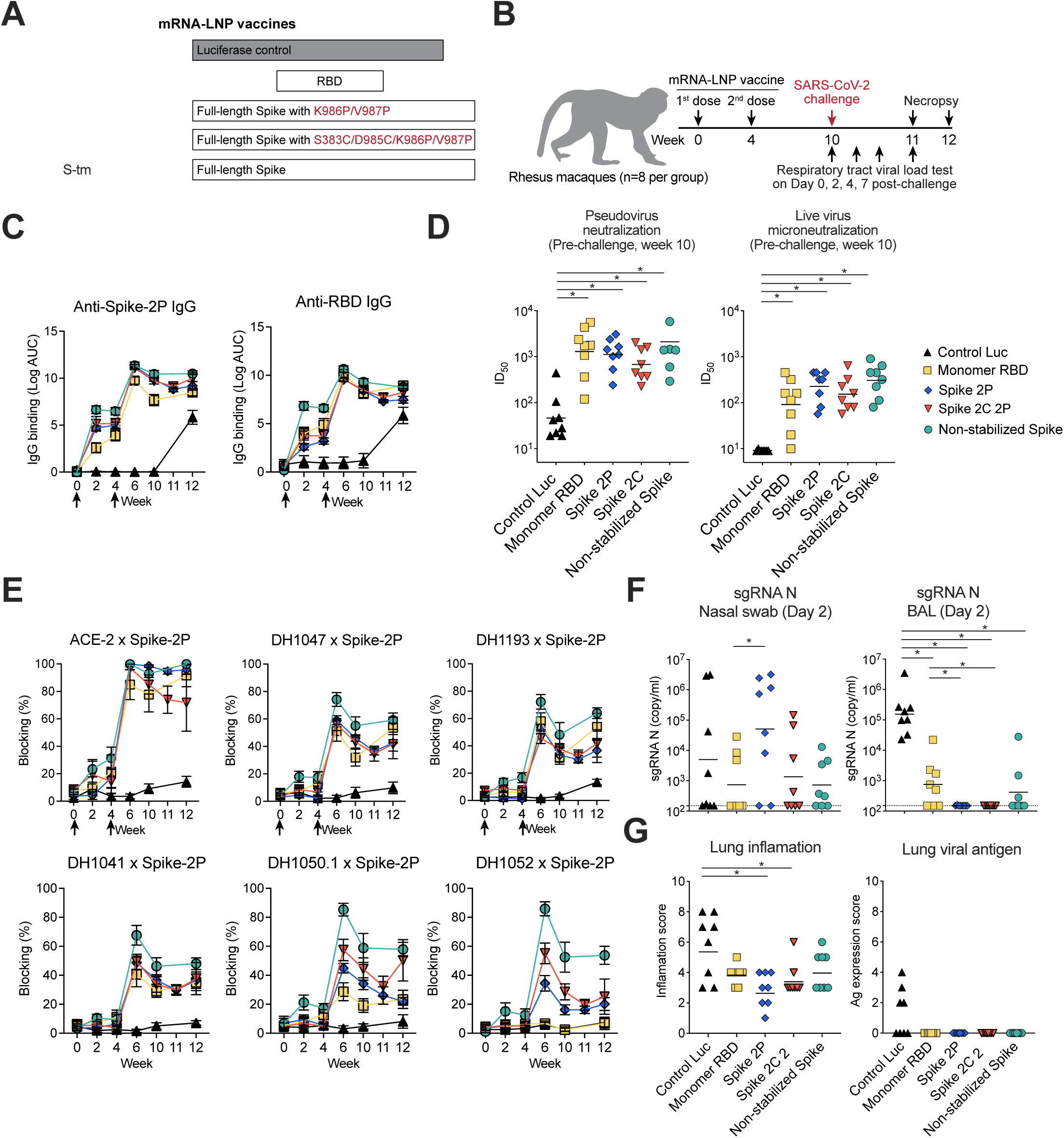
SARS-CoV-2 mRNA-lipid nanoparticle (LNP) vaccines elicited neutralizing antibodies in rhesus macaques. (A) Schematic diagram of the mRNA-LNP vaccines. mRNA-LNP vaccines that encode monomeric receptor-binding domain (RBD), K986P/V987P mutations stabilized full-length spike (Spike 2P), S483C/D985C/K986P/V987P mutations stabilized full-length spike (Spike 2C 2P), or transmembrane spike (S-tm) were compared. A luciferase expressing mRNA-LNP vaccine was made as a control. (B) Rhesus macaque vaccination and challenge regimen. Rhesus macaque (n=8 per group) were immunized intramuscularly by mRNA-LNP vaccine two times in weeks 0 and 4, followed by 10^5^ PFU of SARS-CoV-2 challenge at week 10. Nasal swab and Bronchoalveolar lavage (BAL) samples were collected on post-challenge days 0, 2, 4, and 7. (C) Vaccine-induced SARS-CoV-2 specific IgG binding titers. Serum IgG binding to Spike 2P (S-2P) and RBD was tested by ELISA and shown as log area under the curve (AUC). Symbols indicate the group mean value ± SEM of three replicates. Arrows indicate timing of immunization. (D) SARS-CoV-2 mRNA-lipid nanoparticle (LNP) vaccines elicited neutralizing antibodies against SARS-CoV-2 viruses in rhesus macaques. ID_50_ titers of serum micro- and pseudovirus neutralization of SARS-CoV-2 D614G infection of ACE2-expressing 293T cells. Serum was examined after two immunizations. Each symbol indicates one animal. (E) Sera blocking of ACE-2, RBD neutralizing antibodies DH1041 and DH1047, NTD neutralizing antibodies DH1050.1, and NTD non-neutralizing antibody DH1052 binding to spike. The group mean percentage ± SEM of blocking is shown. Arrows indicate timing of immunization. (F) Reduced SARS-CoV-2 viral replication in the lower respiratory tract of vaccinated macaques. SARS-CoV-2 nucleocapsid gene (N gene) sgRNA was quantified in bronchoalveolar lavage (BAL) and nasal swab samples on Day 2. (G) Quantification of lung inflammation and lung viral nucleocapsid positivity determined by immunohistochemistry of lung tissue sections seven days after challenge. In each panel, symbols represent individual macaques.

### Spike vaccination protected against SARS-CoV-2 challenge in rhesus macaques

To compare the protective efficacy of mRNA-LNP vaccines, we challenged macaques vaccinated with either the S-tm, the S-2P or the S-2C,2P mRNA-LNP with SARS-CoV-2 WA-1/2020 via intratracheal and intranasal routes six weeks after the second immunization (**Fig. 1B**). We quantified SARS-CoV-2 replication in bronchoalveolar lavage (BAL) fluid and nasal swab samples by measuring SARS-CoV-2 envelope (E) and nucleocapsid (N) gene subgenomic RNA (sgRNA). Two and four days following challenge, levels of E and N gene sgRNA in BAL samples from macaques administered each of the spike or RBD vaccines were significantly decreased as compared to Luc mRNA-LNP-immunized macaques (Exact Wilcoxon *P*<0.05 n = 8 macaques) (**Fig. 1F and S3**). Macaques administered the S-2P mRNA-LNP were the only macaques to exhibit undetectable E and N gene sgRNA on day 4 and 7 in BAL fluid, while macaques in other groups had detectable BAL E and N gene sgRNA (**fig. S3**). Immunohistochemistry detected N antigen in the lungs of most Luc mRNA-LNP-immunized macaques on day 7 post-infection (**Fig. 1G)** with some macaques showing undetectable N antigen on day 7 (**fig. S3**). In contrast, N antigen was not detected in the lungs in either S or RBD-immunized macaques (**Figs. 1G and S3B**). Haematoxylin and eosin staining seven days post-challenge showed Luc mRNA-LNP-immunized macaques had the highest inflammation score (**Fig. 1G)**. Thus, S-2P, S-tm, and RBD mRNA-LNP vaccine-mediated immunity each controlled SARS-CoV-2 lung virus replication and lung pathology caused by infection.

In contrast to the lower airway, E and N sgRNA concentrations in nasal swab fluid for all S or RBD-vaccinated groups were not significantly different from the Luc control group. (Exact Wilcoxon *P*>0.05) (**Fig. 1F and S3**). Plasma IgG binding magnitude across all spike or RBD groups did not increase after SARS-CoV-2 challenge, hence viral antigen from replication was not sufficient to boost serum IgG responses (**Fig. 1C**). Thus, each Spike-based mRNA-LNP vaccine provided protective immunity for control of lower-airway SARS-CoV-2 replication but had minimal effects on upper respiratory tract sgRNA levels.

### S-tm mRNA-LNP vaccination elicited cross-Sarbecovirus nAbs

Given the increased magnitude of plasma blocking of Sarbecovirus and SARS-CoV-2 nAbs in S-tm-immunized macaques(**Fig. 1E**), we isolated monoclonal antibodies from three S-tm mRNA-LNP-vaccinated macaques (**fig. S4**). We sorted single memory B cells that bound to full-length SARS-CoV-2 spike protein and/or RBD (**fig. S4**). From the sorted B cells, 88 paired V_H_ and V_L_ immunoglobulin genes were sequenced. The distribution of V_H_, V_D_, and V_D_ gene segment usage, mutation frequencies, and CDR3 lengths recapitulated the expected distributions typically seen in the macaque repertoire (**Fig. S5**) (*34*). Each antibody was expressed in small-scale and assessed for binding to a panel of SARS-CoV-2, SARS-CoV-1, MERS-CoV, endemic cold coronavirus, and zoonotic coronavirus antigens in ELISA (**Fig. 2A, S6**). We selected 57 SARS-CoV-2-specific antibodies based on their breadth and potency of CoV S or RBD binding for further analysis (**Figs. 2A, S6 and S7**). Eighteen of these antibodies bound to the RBDs of various subsets of SARS-CoV-1, SARS-related bat CoV (RaTG13, RsSHC014, HKU3-1), and SARS-CoV-2-related pangolin CoV (GXP4L) (**Fig. 2**). More than half of the RBD antibodies bound to both Wuhan-Hu1 and BA.5 Spike proteins (**Fig. 2**). None of the RBD-specific antibodies bound to human endemic coronavirus (OC43, HKU1, 220E, NL63) or MERS-CoV S antigens (**Fig. 2A and S7**). Among the remaining 39 antibodies, twelve were SARS-CoV-2-specific RBD antibodies (**fig. S7**). Fifteen were NTD-directed, with eleven of these antibodies exhibiting cross-reactivity with animal zoonotic coronavirus isolates GXP4L, RsSHC014, and RaTG13 (**fig. S7**). Twelve of the selected antibodies bound to an unidentified epitope outside of the RBD and NTD and cross-reacted with at least one animal coronavirus spike protein (**fig. S7**).

**Figure 2.**
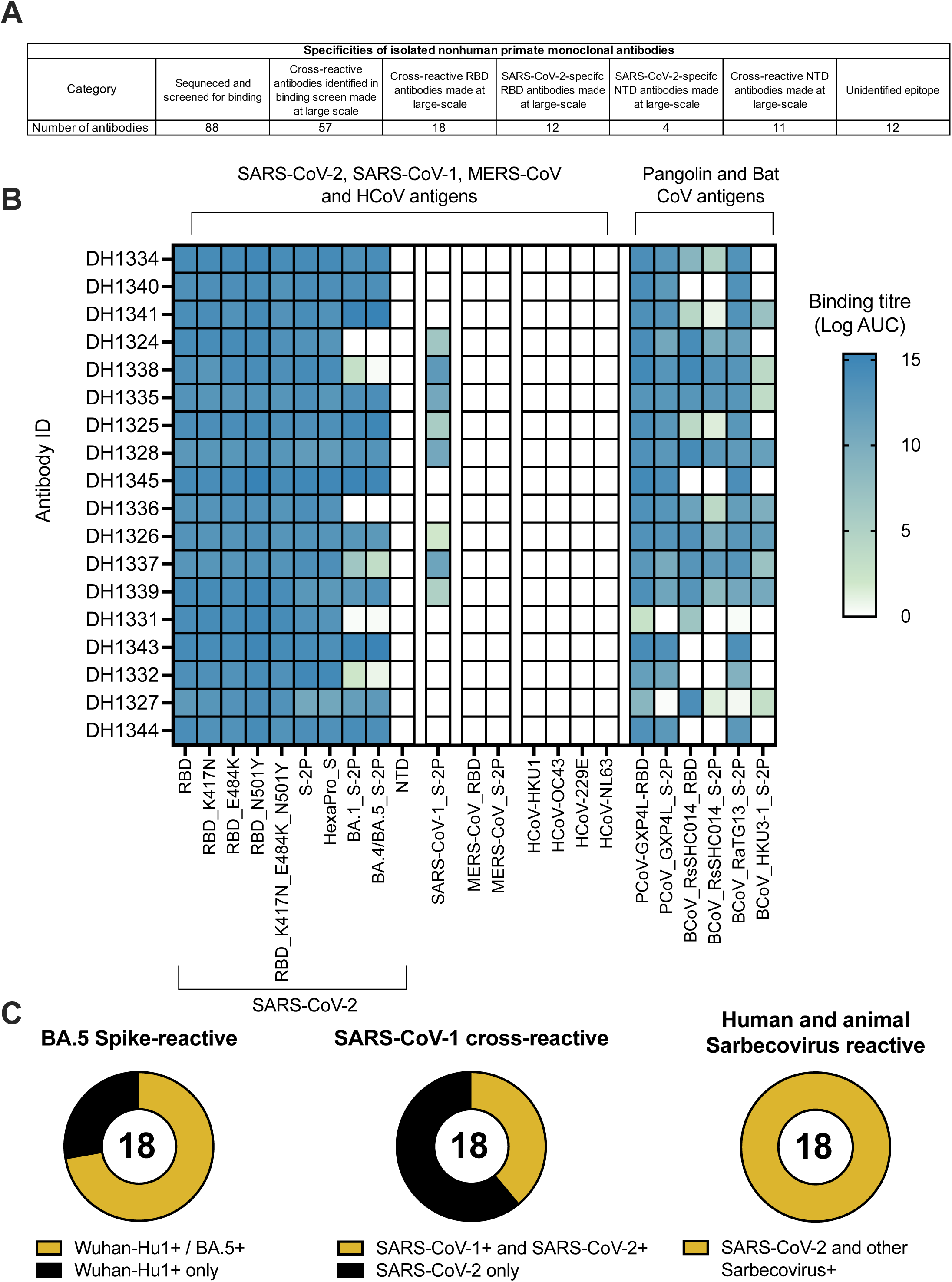
Monoclonal antibodies isolated from wild-type mRNA-LNP immunized macaques cross-react with SARS-CoV-2 variants and SARS-related betacoronaviruses. (A) Summary table of specificities of isolated macaque monoclonal antibodies. (B) Monoclonal antibodies isolated from S-tm mRNA-LNP vaccinated rhesus macaques bind to ectodomain, RBD, NTD from SARS-CoV-2, SARS-CoV-1, SARS-related animal betacoronaviruses, MERS-CoV, and endemic human coronaviruses. The mean log AUC values are shown in the heatmap. (C) Proportion of BA.5 spike reactive, SARS-CoV-1 cross reactive, and human and animal Sarbecovirus-reactive monoclonal antibodies isolated from wild-type mRNA-LNP immunized macaques.

### Vaccine-induced broadly neutralizing monoclonal Abs that targeted the RBD outer face

Next, we performed competitive binding ELISA assays with a subset of 22 cross-reactive or SARS-CoV-2-specific vaccine-induced RBD mAbs and structurally-defined human CoV nAbs. We identified 7 RBD antibodies that blocked >50% of RBD outer face nAbs DH1044 and/or DH1193 (34) binding at ng/mL concentrations (**Fig. 3A and S8A).** Two of these outer face antibodies (DH1329and DH1333) blocked SP1-77, a potent neutralizing antibody against SARS-CoV-2 VOCs (*35*). All seven rhesus antibodies that blocked DH1044 or DH1193 were broad neutralizers of SARS-CoV-2 VOCs, and DH1341, DH1343, DH1344, and DH1345 neutralized all tested SARS-CoV-2 VOCs including Omicron sublineage pseudoviruses (**Fig. 3B-D, and S8**). Three additional antibodies blocked DH1044, but were minimally neutralizing or non-neutralizing (**fig. S8**). Similar to DH1044, no outer face antibodies exhibited cross-neutralization of SHC014 or SARS-CoV-1 (**Fig. 3B**). Finally, we confirmed the epitopes of the outer face nAbs by performing negative stain electron microscopy of the antigen binding fragments (Fabs) or the full-length IgG in complex with SARS-CoV-2 S-2P. The SARS-CoV-2 nAbs DH1333, DH1329, and DH1345 bound to the outer face of the RBD (**Fig. 4A)**, explaining their competition with DH1044 or DH1193 for binding to spike (**Fig. 3A**). DH1333 and DH1329 bound down RBDs with a binding mode similar to DH1044 (**Fig. 4B**); whereas DH1345 bound up RBDs at a site most similar to the RBD-4 community epitope recognized by DH1193 (**Fig. 4C and S2**).

**Figure 3.**
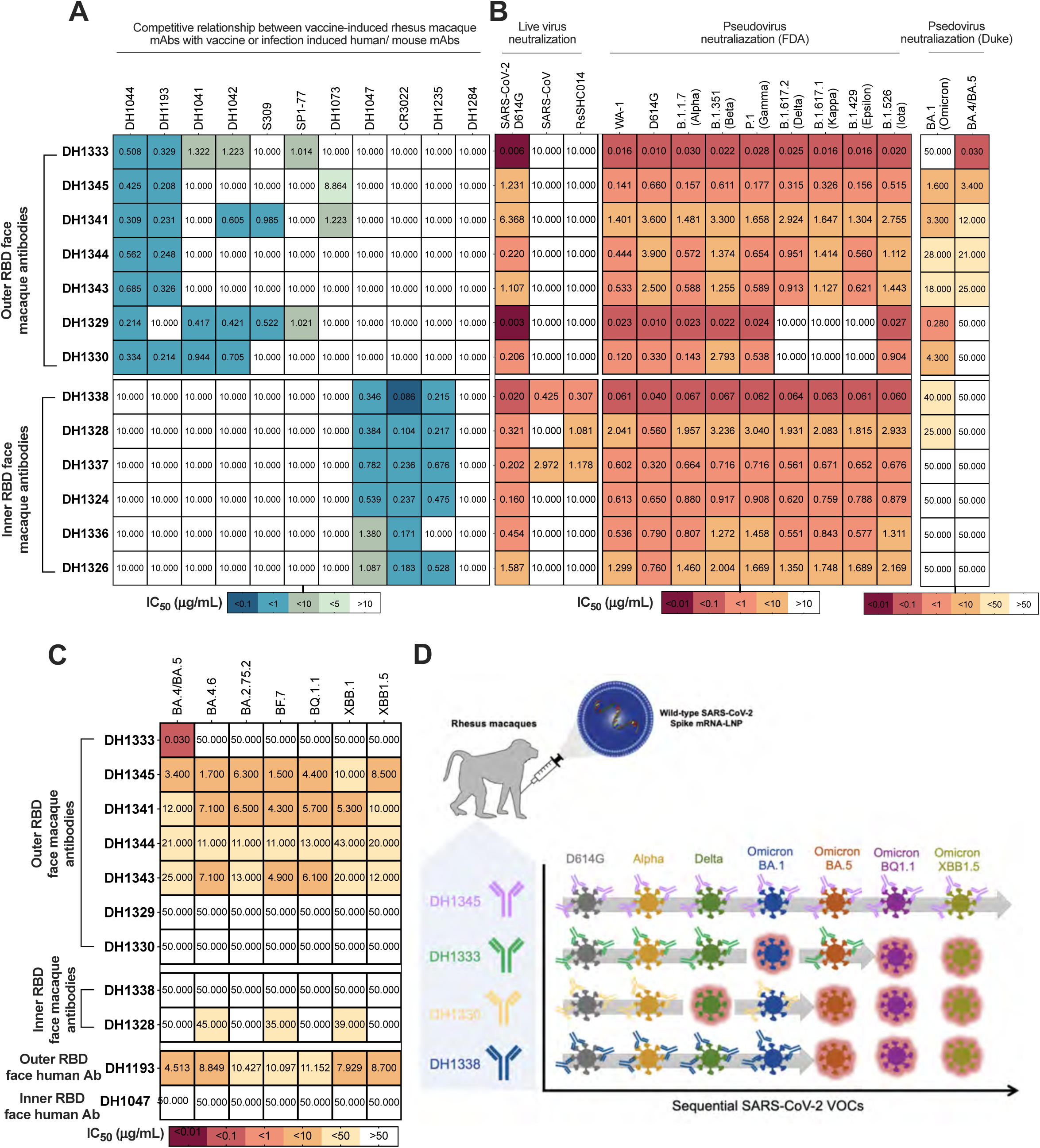
Distinct antibody clones confer pan-SARS-CoV-2 variants of concern neutralization or Sarbecovirus cross-neutralization. (A) The IC50 titer for vaccine-induced rhesus macaque cross-neutralizing RBD antibodies blocking potent vaccine-induced or infection-induced human RBD-reactive monoclonal antibodies. (B) Monoclonal antibodies isolated from S-tm mRNA-LNP vaccinated rhesus macaques neutralize replicating SARS-CoV-2, SARS-CoV, RsSCHC014 viruses, and SARS-CoV-2 pseudovirus variants of concern. (C) Monoclonal antibody neutralization against pseudoviruses of SARS-CoV-2 Omicron sublineages (BA.4/BA.5, BA.4.6, BA2.75.2, BF.7, BQ.1.1, XBB.1 AND XBB1.5) variants in 293T-ACE2 cells. (D) Summary of monoclonal nAb activity against various SARS-CoV-2 VOCs. Rows show neutralization of sequential SARS-CoV-2 VOCs by four individual clones. Columns represent neutralization against the virus listed at the top of the column. Lack of neutralization is shown as a break in the gray arrow and outlining the virus in salmon.

**Figure 4.**
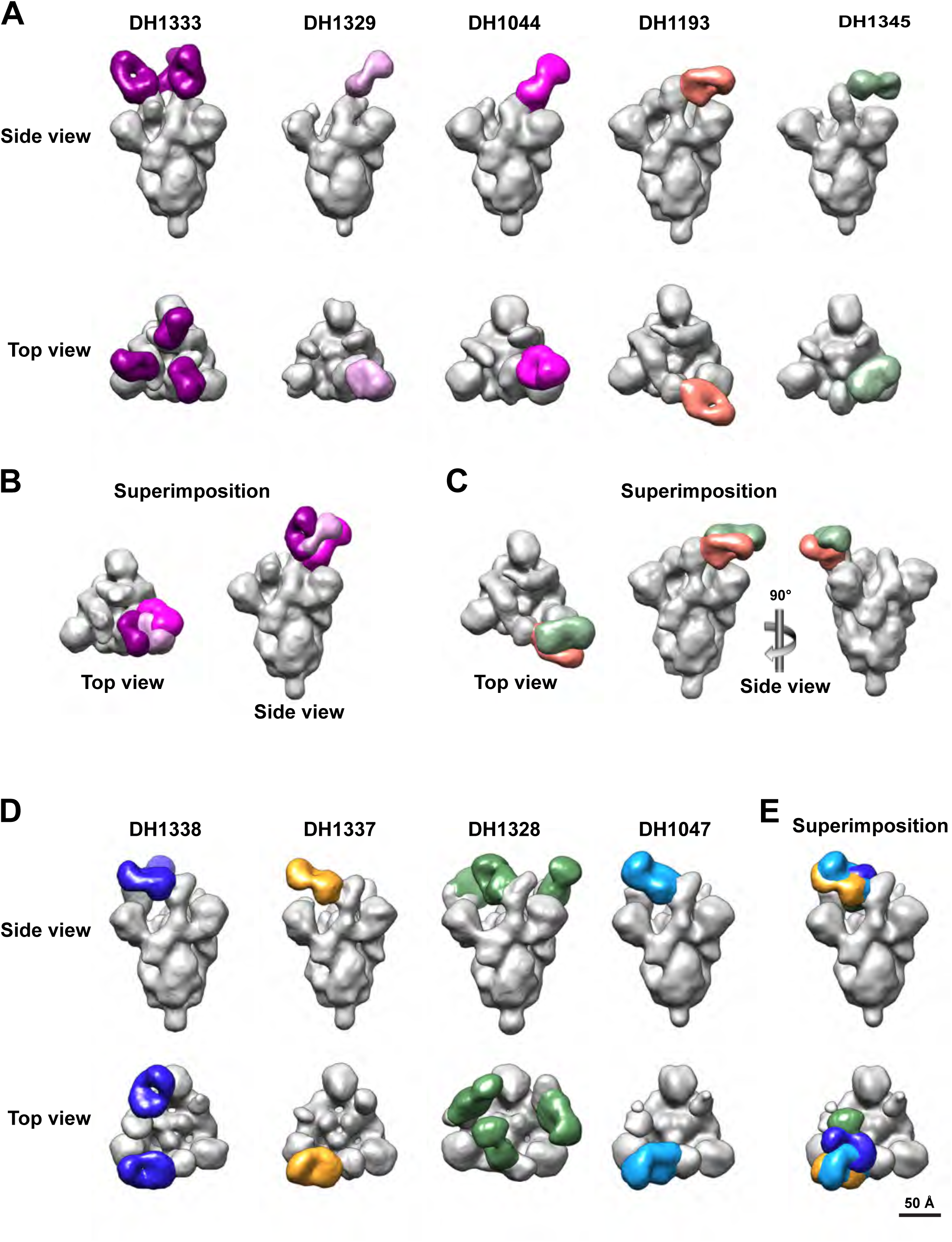
Structural similarities of vaccine-induced neutralizing macaque antibodies and infection-induced human neutralizing antibodies. Structure of antibody bound to spike was determined by negative stain electron microscopy. 3D reconstruction of vaccine-induced and infection-induced cross-neutralizing antibodies (various colors) bound to SARS-CoV-2 Spike (gray). (A) The epitopes of broad SARS-CoV-2-neutralizing RBD antibodies that target outer RBD face. (B-C) Superimposition of Fab regions to compare binding angles of approach for vaccine-induced antibodies (B) DH1329 and DH1333 or (C) DH1345 with infection induced human outer face antibodies DH1044 or DH1193. (D) The epitopes of cross-neutralizing RBD antibodies that target inner RBD face. (E) Superimposition of Fab regions to compare human infection-induced cross-Sarbecovirus neutralizing antibody DH1047 with vaccine-induced antibodies (DH1328, DH1337, DH1338). Fab regions are depicted in different colors and SARS-CoV-2 S-2P spike protein is shown in gray.

### Vaccine-induced cross-sarbecovirus nAbs that targeted the inner face of RBD

Among the subset of 22 cross-reactive or SARS-CoV-2-specific vaccine-induced RBD mAbs tested, six antibodies blocked cross-nAbs DH1047 and DH1235 which bind to the inner face of the RBD and broadly neutralize Sarbecoviruses (*30–32, 36*) (**Fig. 3A**). The six macaque antibodies also blocked CR3022 binding to spike (**Fig. 3A**). CR3022 is a SARS-CoV-1 nAb that cross-reacts with SARS-CoV-2 (*37, 38*). Consistent with targeting the DH1047 inner face epitope, RBD nAbs DH1338 and DH1337 neutralized SARS-CoV-2, SARS-CoV, and bat CoV RsSCH014 (**Fig. 3B**). A third antibody, DH1328 neutralized replication-competent SARS-CoV-2 D614G and BatCoV RsSHC014 (**Fig. 3B**). One additional antibody blocked outer face antibodies, but was non-neutralizing (**fig. S8**). Like DH1047, none of the inner face antibodies neutralized BA.4/5 virus (**Fig. 3C**). Thus, mRNA-LNP S-tm vaccination generated cross-Sarbecovirus nAbs that were distinct from the Omicron sublineage broad nAbs.

Negative stain electron microscopy of the antibody-spike protein complexes confirmed that the cross-Sarbecovirus nAbs (DH1338, DH1337, DH1328) bound to the inner face of RBD similar to DH1047 (**Figs. 4D-E** and **S6D**). DH1328 targeted the same epitope as DH1047 but had a slightly different binding angle of approach compared to DH1338, DH1337, and DH1047 (**Fig. 4D-E**). Thus, NSEM analysis demonstrated that vaccine-induced inner face antibodies bound an epitope overlapping with RBD-6 and7 community epitopes just like human nAb, DH1047.

### Dynamic SARS-CoV-2 VOCs sensitivity to neutralizing monoclonal RBD Abs

Neutralization resistance mutations have accumulated in SARS-CoV-2 since the beginning of the COVID-19 pandemic (*39–45*). Next, we examined how vaccine-induced RBD nAbs neutralized SARS-CoV-2 VOCs that arose at various points during the COVID-19 pandemic. First, six of the nAbs, including DH1338, neutralized 8 of the variants but lacked neutralization of either BA.1 or BA.4/5 (**Fig. 3B and C**). Second, antibodies like DH1329 and DH1330 neutralized early variants, lost neutralization activity against Delta, but also neutralized Omicron BA.1 (**Fig. 3B-D**), which arose after the Delta VOC. Third, DH1333 exhibited highly potent neutralization for all VOCs observed prior to the BA.1 VOC (**Fig. 3B-D**). Although, DH1333 did not neutralize BA.1, it neutralized BA.4/5 pseudovirus with a 30 ng/mL IC50 titer (**Fig. 3B-D**). Thus, the vaccine induced a polyclonal RBD neutralizing antibody response had multiple neutralization patterns against VOCs.

### Structural analysis of inner RBD human and macaque broad Sarbecovirus neutralizing antibodies

Next, to visualize the DH1338 epitope, we used cryo-EM to determine a structure of the DH1338 antigen-binding fragment (Fab) bound to the SARS-CoV-2 S to an overall resolution of 3.1 Å and performed local refinement to further resolve the Fab-RBD interface at 3.4 Å **(Fig. 5A-E, S9, table S1-2).** Like DH1047, the epitope of DH1338 is on the RBD inner face and overlaps with the ACE2 binding site on the RBD (**Fig. 5E**). The RBD-epitope of DH1338 resembled that of S2X259, and overlapped with the DH1047 epitope, although DH1338 had a smaller binding footprint compared to DH1047 (**Fig. 5B-E, S10**). Like S2X259 and DH1047, DH1338 utilized its HCDR3 loop to contact the RBD β −2 strand (**Fig. 5C**). The DH1338 HCDR3 residue Tyr100D makes extensive H-bonds with residues Ser371, Thr376, Phe377, and Tyr369 in the RBD ß-2 strand. The light chain buried surface area (BSA) at the RBD interface is dominated by LCDR1 interactions (**Table S1-2**). Four residues in the LCDR1 Ser28, Ser30, Ser31, and Tyr32, as well as residue Ser67 in the framework region contact the RBD . The LCDR1 residues Ser30 and Ser 31 H-bond via their side chain hydroxyls with the side chain of residue Asp405, and Tyr32 H-bonds via its side chain hydroxyl with the carbonyl oxygen of RBD residue Gly404 and the side chain of residue Asp405, respectively. Another H-bond was observed between the main chain carbonyl oxygen of the LCDR3 residue Ser92 and main chain nitrogen of RBD residue Val503. Hence, the non-stabilized S mRNA-LNP-vaccinated macaques elicited cross-nAbs that used different molecular interactions to bind to the same RBD epitope as DH1047.

**Figure 5.**
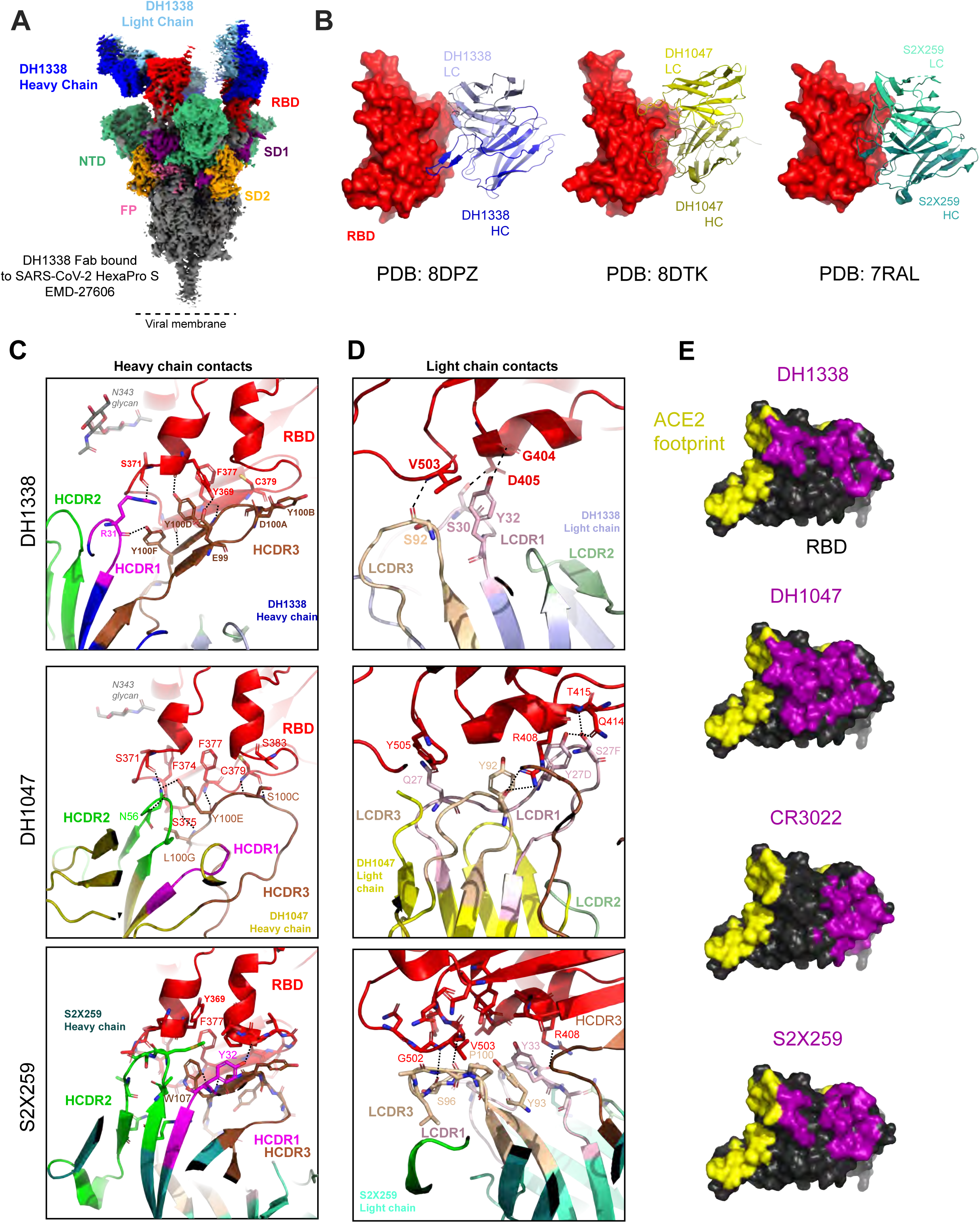
Cryo-EM reconstruction of DH1338 Fab bound to SARS-CoV-2 S protein. (A) 3-Fab bound complex of DH1338 to SARS-CoV-2 S HexaPro Protein reconstruction (EMD-27606). Colored by region. (B) Antibody binding interface. RBD shown in surface representation and colored red, with DH1338 shown in blues, DH1047 shown in yellows, and S2X259 shown in teals. (C) Top, DH1338 HCDR3 loop shown in brown, polar contacts with RBD, shown in black dashed lines. Middle, DH1047 HC polar contacts with RBD. Bottom, S2X259 HC polar contact with RBD. (D) Light chain polar contacts of DH1338 (top), DH1047 (middle), and S2X259 (bottom). (E) RBD is shown in black and ACE2 binding footprint is colored yellow. Binding footprints of antibodies DH1338 (PDB: 8DPZ), DH1047 (PDB: 8DTK), CR3022 (PDB: 6YLA), and S2X259 (PDB: 7RAL) are shown in purple. All RBDs were aligned to PDB: 6M0J, for RBD representation.

### Vaccine induction of protective antibodies against a pre-emergent BatCoV

DH1338 exhibited broad neutralization of Sarbecoviruses (**Fig. 3**), thus we investigated its ability to protect against a mouse-adapted bat Sarbecovirus, RsSHC014-MA15. RsSHC014 is a SARS-like virus from Chinese horseshoe bats with potential for human infection since it can infect human primary airway cells and binds human ACE2 (*29*). We challenged mice with 1×10^4^ PFU of pathogenic RsSHC014 MA15 virus (**Fig. 6A**). Twelve-month-old BALB/c mice were passively administered 300 μg of DH1338 12 hours prior to infection as prophylaxis or 12 hours post-infection as a therapy (**Fig. 6A**). Negative control mice received anti-influenza antibody CH65 12 hours prior to infection. Prophylactic DH1338 treatment protected mice from weight loss through 4 dpi and therapeutic treatment halted weight loss at day 2 post infection (**Fig. 6B**). Prophylactic DH1338 administration suppressed infectious virus in the lungs to undetectable levels in all but one mouse (**Fig. 6B**). Therapeutic administration of DH1338 suppressed infectious virus in the lungs to undetectable levels in four of ten mice and lowered infectious virus in the remaining six mice compared to the negative control group (**Fig. 6C**). Additionally, prophylactic administration of DH1338 completely protected all mice from macroscopic lung damage, as determined by the lung discoloration score (**Fig. 6D**). Finally, there was no mortality observed in any of the antibody groups (**Fig. 6E**). These results indicated mRNA vaccination with spike lacking 2P substitutions induced nAbs that provide prophylactic and therapeutic benefits against pre-emergent zoonotic Sarbecovirus infection in an aged mouse model.

**Figure 6.**
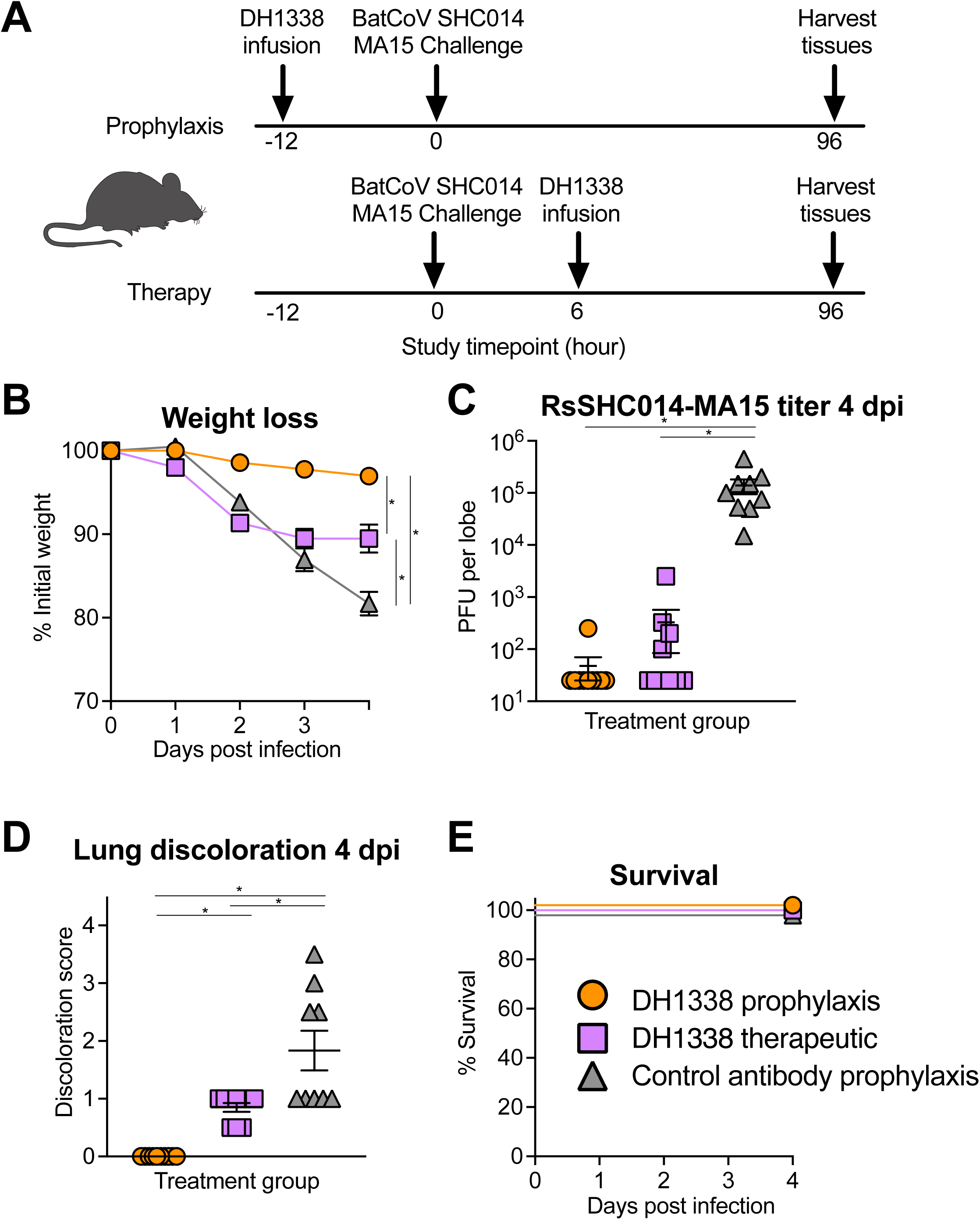
Macaque antibody DH1338 confers prophylactic and therapeutic protection against batCoV RsSHC014 replication in aged mice. (A) Scheme of prophylactic and therapeutic administration of DH1338 in mice. DH1338 or control antibody, CH65, were administered either prophylactically 12 hours before infection, or therapeutically 12 hours after infection. Disease-susceptible aged mice (n=10) were challenged with a mouse-adapted bat coronavirus, SHC014-MA15. (B) The percent weight loss is shown for mice treated prophylactically or therapeutically. Values shown are group mean ± SEM. (C) Lung viral replication of SHC014-MA15 is shown for mice treated prophylactically or therapeutically with DH1338 and CH65 at 4 days post-infection (dpi). (D) Lung discoloration scores are shown for mice treated with DH1338 or CH65 prophylactically and therapeutically. (E) Percent survival is shown for all three groups challenged with SHC014-MA15. Horizontal bars are mean± SEM. *, Exact Wilcoxon *P*<0.05 n = 8-10 mice.

## DISCUSSION

Here, we show that the Wuhan-Hu-1 spike delivered as an mRNA can induce broadly reactive antibodies that either bind to the RBD outer face and neutralize multiple late Omicron variants or bind to the RBD inner face and neutralize SARS-CoV-1 and preemergent Sarbecoviruses. The 2P substitutions were not required for induction of such neutralizing antibodies. Moreover, we demonstrate different neutralization patterns by the Wuhan-Hu-1 RBD-induced antibodies. A subset of antibodies were capable of neutralizing XBB.1.5 despite being elicited by the Wuhan-Hu-1 ancestral isolate of SARS-CoV-2. Thus, these results show antibodies induced by ancestral versions of SARS-CoV-2 Spike have the potential to contribute to neutralization of current SARS-CoV-2 isolates. These results also show that there is a neutralizing antibody epitope that remains conserved between XBB.1.5 and the ancestral isolates of SARS-CoV-2. Our structural studies identified the DH1193 epitope as one of these conserved epitopes on SARS-CoV-2 S variants targeted by macaque antibodies. Thus, both human infection-induced and macaque vaccine-induced antibodies can target either the outer or the inner faces of the RBD to broadly neutralize CoVs (*30–32, 36*).

The neutralization patterns of the broadly neutralizing Sarbecovirus antibodies showed that Omicron sublineages have mutated the inner face epitope that is conserved on SARS-CoV-1 and animal Sarbecoviruses. Hence, vaccines aiming to elicit the broadest response against both SARS-CoV-2 VOCs and SARS-like Sarbecoviruses should aim to elicit antibodies to both RBD inner and outer face epitopes. Wuhan-Hu-1 S-tm mRNA-LNP vaccines here and S nanoparticle vaccines in previous studies (*46, 47*) elicited both responses in nonhuman primates, suggesting next-generation vaccines may only need to selectively boost antibodies targeting the most conserved epitopes on each face of the RBD. Designing resurfaced receptor binding domains that present only the conserved epitopes in the outer and inner faces would be one immunogen approach to directing antibodies to these conserved epitopes. Such vaccine approaches may be necessary since we observed only four omicron sublineage nAbs out of 88 total antibodies.

In summary, pan-coronavirus vaccines may benefit from the contributions of distinct broadly neutralizing antibodies that cooperate to provide immunity against SARS-CoV-2 VOCs and other Sarbecoviruses. Our findings show that conserved neutralizing epitopes present on ancestral SARS-CoV-2 RBD are present on the outer face of SARS-CoV-2 VOCs including XBB.1.5 RBD, as well as the inner face of pre-emergent Sarbecovirus RBDs. Boosting regimens will need to target these two epitopes to increase their immunogenicity and to convert such antibody responses to dominance. Immunogen designs such as non-stabilized spikes or engineered RBD nanoparticles (*48*) may be strategies that can convert subdominant antibody responses to dominant responses. Finally, strategies of full-length S should consider non-stabilized S-tm as vaccine candidates.

## MATERIALS AND METHODS

### Rhesus macaques and immunizations

Rhesus macaques used in this study were housed and treated in AAALAC-accredited institutions. The study protocol and all veterinarian procedures were approved by the Bioqual IACUC per a memorandum of understanding with the Duke IACUC and were performed based on standard operating procedures. Nucleoside-modified messenger RNA encapsulated in lipid nanoparticles (mRNA-LNP) encoding different versions of SARS-CoV-2 spike or RBD was prepared as previously stated(*33*). Rhesus macaques were on average 8 or 9 years old, and their body weights ranged from 2.75 to 8 kg. 8 rhesus macaques were allocated into each group and immunized intramuscularly with 50µg of mRNA-LNP encoding the monomer receptor-binding domain (RBD), full-length spike protein stabilized with K986P/V987P mutations (Spike 2P), full-length spike protein stabilized with S483C/D985C/K986P/V987P (Spike 2C 2P), or wild-type spike without any stabilization mutations. Balance of male and female animals per group was considered when macaque availability permitted. Studies were performed unblinded. Animals were observed and evaluated by Bioqual veterinary staff throughout immunizations and no significant abnormalities were noted in obtained CBCs and chemistries.

### SARS-CoV-2 intranasal and intratracheal challenge

Infectious dose of 10^5^ PFU of SARS-CoV-2 (2019-nCoV/USA-WA1/2020) was selected to challenge all animals. Aliquots of virus were stored at −80°C and thawed by hand and placed immediately on wet ice. Prior to inoculation, the stock was diluted to 2.5×10^4^ PFU/mL in PBS and vortexed for 5 seconds and 3mL of diluted virus were administered to animals in combined intratracheal and intranasal routes. Nasal swabs, bronchoalveolar lavage (BAL), plasma, and serum samples were collected at 0, 2, 4, 7 days post-challenge. Animals were necropsied and lungs were harvested on Day 7 post-challenge for histopathology and immunohistochemistry (IHC) analysis.

### SARS-CoV-2 pseudovirus neutralization

Neutralization of SARS-CoV-2 Omicron Spike-pseudotyped virus was performed by adopting an infection assay described previously (*49*). Mutations for BA.1, BA.4/5, BA.4.6, BA.2.75.2, BF.7, BQ.1., XBB.1, or XBB.1.5 were introduced into an expression plasmid encoding codon-optimized full-length spike of the Wuhan-1 isolate (VRC7480) VRC7480 by site-directed mutagenesis and confirmed by full-length spike gene sequencing by Sanger Sequencing. Pseudovirions were produced in HEK293T/17 cells following a co-transfection of spike plasmid, lentiviral backbone plasmid (pCMV ΔR8.2) and firefly Luc reporter gene plasmid (pHR’ CMV Luc) in a 1:17:17 ratio in Opti-MEM. Transfection mixtures were then added to pre-seeded HEK 293 T/17 cells and incubated for 16–20 hours at 37°C. After the incubation, medium was removed and replaced with fresh growth medium. A pre-determined dose of virus was incubated with serially diluted monoclonal antibodies or heat inactivated serum. HEK 293T/17 cells expressing ACE2 were suspended using TrypLE express enzyme solution and immediately added to all wells. Control wells received HEK 293T/17 cells and virus control and another set received cells only and was assessed as background control. After 66–72 h of incubation of cells and monoclonal antibodies, medium was removed, 1x lysis was added and incubated for 10 minutes at room temperature. At the end of the incubation, Bright-Glo luciferase reagent was added to all wells. After 1–2 minutes, luminescence was measured using a luminometer. Neutralization titers were quantified as serum dilution (ID50/ ID80) at which relative luminescence units (RLU) were reduced by 50% and 80% compared to virus control wells after subtraction of background.

For pseudovirus assays with SARS-CoV-2 WA-1, D614G, B1.1.7, B.1.351, P.1, B.617.2, B.617.1, B.1.429, B.1.526, 50 µL of SARS-CoV-2 S pseudovirions were pre-incubated with an equal volume of medium containing serum at varying dilutions at room temperature for 1 h, then virus-antibody mixtures were added to 293T-ACE2 (WA-1 and B.1.351 assays) or 293-ACE2-TMPRSS2 (WA-1 and P.1 assays) cells in a 96-well plate. After a 3 h incubation, the inoculum was replaced with fresh medium. Cells were lysed 24 h later, and luciferase activity was measured using luciferin. Controls included cell only control, virus without any antibody control and positive control sera. Neutralization titers are the serum dilution (ID50/ID80) at which relative luminescence units (RLU) were reduced by 50% and 80% compared to virus control wells after subtraction of background RLUs.

### Live virus neutralization assays

Live virus assays were performed with SARS-CoV-1 Urbani, SARS-CoV-2 D614G, and RsSHC014 designed to express nanoluciferase (nLuc) (*50, 51*). For the live virus neutralization assays, cells were plated at 20,000 cells per well in clear bottom, black-walled 96-well plates the day prior to the assay. On the day of the assay, monoclonal antibodies were serially diluted 3-fold for eight dilutions. Serially diluted antibody was added at a 1:1 volume with diluted virus and incubated for 1hr. Antibody-virus dilutions were then added to cells and incubated at 37°C with 5% CO2. Following a 24hr incubation, luminescence was generated by adding 25 μL of Nano-Glo Luciferase Assay System (Promega). Spectramax M3 plate readers (Molecular Devices) were used to measure luminescence. The inhibitory concentration that resulted in a fifty percent reduction in luminescence virus was calculated using GraphPad Prism via four-parameter dose-response curves.

### Serum and antibody binding ELISA

For direct binding assays, 384-well plates were coated with 15µl/well of diluted antigen in 0.1 M sodium bicarbonate and stored at 4°C overnight. On the following day, plates were washed with PBS +0.05% Tween 20 (Superwash) and subsequently blocked with 40µl/well using PBS containing 4% (w/v) whey protein, 15% Normal Goat Serum, 0.5% Tween-20, and 0.05% Sodium Azide (SuperBlock) for 1 hour at room temperature. Purified monoclonal antibodies were serially diluted (3-fold). Positive and negative controls for each antigen were included for each plate. The starting concentration of the purified monoclonal antibodies was typically 100µg/ml and for serum 1:30. After 1 hour of blocking, plates were washed with Superwash and serially diluted samples and controls were added to the plate at 10µl/well and incubated plates for 1 hour at room temperature. After the incubation, plates were washed with Superwash and 10 µl/well of HRP-conjugated secondary antibody diluted in Superblock without Sodium Azide was added to the plates and incubated for 1 hour at room temperature.

After incubation, plates were washed with Superwash and TMB substrate was added at 20µl/well and incubated at room temperature for 15 minutes. After the incubation, the reaction was stopped using 1% HCl solution at 20µl/well. Absorbance values were read at 450nm using SoftmaxPro ELISA microplate reader.

### ACE2 and neutralizing antibody blocking assay

Recombinant ACE-2 protein was diluted to 2 µg/mL in filtered 0.1M sodium bicarbonate. 384 well ELISA plates were coated with 15 µL/well of coating solution and incubated overnight at 4°C. Following day, plates were washed and blocked with 40 µL/well using 3% BSA in PBS. Plates were then incubated for 1 hour at room temperature. In a separate tube, SARS-CoV-2 spike was diluted to 0.4 µg/ml in 1% BSA/0.05% Tween-20 in PBS. Macaque sera were diluted to 1:25. If purified antibodies were used, the serial dilution of antibodies was performed and the starting concentration was 100 µg/ml, titrated 3-fold in 1% BSA/0.05% Tween-20 in PBS. Spike-2P protein was mixed with the diluted sera or antibodies at a final Spike concentration equal to the EC50 at which spike binds to ACE-2 protein. The mixture was incubated at room temperature for 1 hour.

After the blocking step, plates were washed, and the antibody-spike mixture was added to ACE2 coated wells and incubated for 1 hour at room temperature. Plates were washed and a polyclonal rabbit serum against SARS-CoV-2 spike was added and incubated for 1 hour following a wash step and detection of binding with goat anti rabbit-HRP (Abcam catalog # ab97080). TMB substrate was added to the plates for detection of HRP. Hydrochloric acid was added to the plat to stop the detection reaction. Well absorbance was read at 450 nm. Percent blocking was calculated as follows: blocking % = (100 − (Optical Density of antibody plus Spike/Optical Density of Spike only)*100).

### Plasma and IgG blocking of RBD monoclonal antibody binding

Blocking assays for DH104, DH1042, DH1044, DH1047, DH1073, DH1235, DH1284, DH1193, DH50.1, DH1052, S409, CR3022, SP1-77 were performed as stated above for ACE2, except plates were coated with the binding antibody of interest instead of ACE2 and used biotinylated version of the binding antibody as detection antibody.

### Subgenomic RNA real time PCR quantification

The measuring of SARS-CoV-2 E gene and N gene subgenomic mRNA was done using one-step RT-qPCR. The assay was adapted from previously described methods as described previously (49).

### Histopathology and immunohistochemistry

Lung specimen from individual animals were fixed in 10% neutral-buffered formalin, processed, and blocked in paraffin for histology analyses. All tissues were sectioned at 5 µm and stained with hematoxylin-eosin (H&E) to assess histopathology. Stained sections were evaluated by a board-certified veterinary pathologist in a blinded manner. Olympus BX51 microscope and Olympus DP73 camera were used to obtain the photographs. The representative images for each group are shown.

For immunohistochemistry (IHC) staining for SARS-CoV-2 nucleocapsid antigen, Bond RX automated system with the Polymer Define Detection System (Leica) was used following the manufacturer’s protocol. Briefly, tissue sections were dewaxed with Bond Dewaxing Solution (Leica) at 72°C for 30 minutes. At the end of incubation, the sections were rehydrated with graded alcohol washes and 1x Immuno Wash (StatLab). Heat-induced epitope retrieval (HIER) was performed using Epitope Retrieval Solution 1 (Leica) and along with heating the tissue section to 100°C for 20 minutes. To quench endogenous peroxidase activity prior to applying the SARS-CoV-2 nucleocapsid antibody, a peroxide block (Leica) was applied for 5 min (1:2000, GeneTex, GTX135357). To minimize the background, antibodies were diluted in Background Reducing Antibody Diluent (Agilent) and subsequently incubated with an anti-rabbit HRP polymer (Leica) and colorized with 3,3’-Diaminobenzidine (DAB) chromogen for 10 minutes, and at the end slides were counterstained with hematoxylin.

### Antigen-specific single B cell sorting

Antigen-specific single macaque B cell sorting was performed as described previously with modifications (*52*). Briefly, cryopreserved peripheral blood mononuclear cells (PBMC) were processed and stained with natural killer (NK), T, and B cell surface markers and fluorophore-labeled SARS-CoV-2 Spike and RBD proteins. Antibodies used for staining were CD20 fluorescein isothiocyanate (FITC) clone L27 (BD Biosciences Cat No. 347673), CD3 peridinin chlorophyll protein (PerCP) Cy5.5 clone SP34-2 (BD Biosciences Cat No. 552852), IgD phycoerythrin (PE) polyclonal (Southern Biotech Cat No. 2030-09), CD8 PE Texas Red clone 3B5 (Invitrogen Cat No. MHCD0817), IgM PE Cy5 clone G20-127 (BD Biosciences Cat No. 551079), CD16 PE Cy7 clone 3G8 (BD Biosciences Cat No. 557744), Live / Dead Aqua (Invitrogen Cat No. L34957), CD14 brilliant violet (BV) 570 clone M5E2 (BioLegend Cat No. 301832), and CD27 allophycocyanin (APC) Cy7 clone O323 (BioLegend Cat No. 302816). SARS-CoV-2 Spike and/or RBD reactive, live, IgD-single B cells were sorted into 96-well PCR plate where each B cell is separated into a single well that contained cell lysis buffer and 5X first-strand synthesis buffer and subjected to reverse transcription of RNA.

### Antibody gene RT-PCR

Antibody gene RT-PCR was performed as detailed before with modifications (*52*). Briefly, Superscript III (Thermo Fisher Scientific Cat No. 18080044) and immunoglobulin constant region-specific reverse primers were used to reverse transcribe B cell RNA. Two rounds of nested PCR were performed using 5 μL of complementary DNA (cDNA) to amplify immunoglobulin heavy and light chain variable regions and yielding variable regions were confirmed by using 96 well E-gel agarose electrophoresis (Thermo Fisher Scientific). Purified PCR amplicons were then sequenced with forward and reverse primers and contigs of the amplified immunoglobulin sequences were made by the Duke automated sequence analysis pipeline (Duke ASAP). Immunogenetics of rhesus macaque and human immunoglobulin genes were determined with the macaque reference library using Cloanalyst. Simultaneously, to express antibodies in small scall that are required for binding screens, purified PCR amplicon was used for overlapping PCR and generated a linear expression cassette. The expression cassette was transfected into Expi293F cells (Thermo Fisher Scientific, Cat No. A14527) supernatant of the cell culture media were tested for binding to distinct SARS-CoV-2 spike antigens. Based on the positive binding data, antibodies were down selected for gene synthesis. The synthesized heavy and light chains were cloned into gamma, kappa, or lambda expression vectors (GenScript) and subjected to transient transfection of Expi293F cells purify immunoglobulin heavy and light chain plasmids.

### Recombinant antibody expression

Recombinant antibody expression was performed as described elsewhere (*52*). Briefly, Expi293F cells were co-transfected with heavy and light chain plasmid using Expifectamine (Thermo Fisher Scientific, Cat No. A14526) per the manufacturer’s protocol. Cells were harvested and centrifuged post 5 days transfection, and the cell culture supernatant was concentrated and incubated with protein A beads overnight at 4°C. The mixture of protein A beads (Thermo Fisher Scientific) and supernatant was centrifuged. Protein A resin was washed by resuspending it in 25 mL of Tris-buffered saline pipetted into an empty gravity flow column followed by an elution step using 2% glacial acetic acid. The pH was neutralized by adding 1M Tris pH8.0 to a final pH of 6. The eluate was concentrated and buffer exchanged into 25 mM sodium citrate pH6, 150 mM NaCl pH6. The purified IgG monoclonal antibodies were run in SDS-PAGE followed by Coomassie blue staining and western blot to confirm correct molecular weights.

### SARS-CoV-2 protein production

The human and animal coronavirus ectodomain constructs were produced and purified as described previously (*10, 31*). The full-length spike proteins were stabilized by introducing 2 prolines at amino acid positions 986 and 987 (Spike 2P) and transiently transfected in FreeStyle 293-F cells using Turbo293 (SpeedBiosystems) or 293Fectin (ThermoFisher). The constructs contained an HRV3C-cleavage site followed by His and/or streptagII tags for purification purposes. Post 6-days transfection, protein was purified using cell-free culture supernatant by StrepTactin resin (IBA) and by size-exclusion chromatography using Superdex 200 (RBD and NTD) or Superose 6 (S-2P) column (GE Healthcare) in 10 mM Tris pH8, 500 mM NaCl. In some cases, after affinity chromatography, the tag sequences were cleaved by HRV3C digestion prior to size exclusion chromatography. ACE2-Fc was expressed by transient transfection of Freestyle 293-F cells and purified from cell culture supernatant by HiTrap protein A column chromatography followed by Superdex200 size-exclusion chromatography in 10 mM Tris pH8,150 mM NaCl.

### Negative stain electron microscopy

Antibody-spike complexes with DH1329 and DH1333 were made by mixing the corresponding full-length IgG with D614G 2P spike at a 1.5 to 1 molar ratio; Fab-spike complexes with DH1047, DH1193, DH1328, DH1337, and DH1338 were made by mixing the corresponding Fab with Hexapro spike at a 9 to 1 molar ratio; and Fab-spike complex of DH1044 was made by mixing Fab with 2P spike at a 9 to 1 molar ratio. All complexes were incubated for one hour at 37 °C then processed for NSEM. For NSEM, complexes were brought to room temperature and crosslinked by diluting with HBS-buffer containing 20 mM HEPES, 150 mM NaCl, and 8 mM glutaraldehyde, pH 7.4, with sufficient buffer added for a nominal final spike concentration of 0.2 mg/ml. After crosslinking for five minutes at room temperature, excess glutaraldehyde was quenched by adding sufficient 1 M Tris stock, pH 7.4 for a final Tris concentration of 80 mM and incubating for 5 minutes at room temperature. Quenched samples were then applied directly to a glow-discharged carbon-coated EM grid for 10-12 second, then blotted and stained with 2 g/dL uranyl formate for 1 min, blotted and air-dried. Grids were examined on a Philips EM420 electron microscope operating at 120 kV. Dataset were collected for DH1044 and DH1047 at a nominal magnification of 82,000x and recorded on a 4 Mpix CCD at 4.02 Å/pixel. All subsequent datasets were collected at a nominal magnification of 49,000x and recorded on a 76 Mpix CCD camera at 2.4 Å/pixel. Images were analyzed by 2D class averages, 3D classifications, and final 3D reconstructions calculated using standard protocols with Relion 3.0 (69).

### Cryo-Electron microscopy of antibody-Spike complexes

Purified SARS-CoV-2 spike ectodomains were diluted to a concentration of ∼1.5 mg/mL in 2 mM Tris pH 8.0, 200 mM NaCl and 0.02% NaN3 and 0.5% glycerol was added. A 2.4-μL drop of protein was deposited on a Quantifoil-1.2/1.3 grid (Electron Microscopy Sciences, PA) that had been glow discharged for 10 s using a PELCO easiGlow Glow Discharge Cleaning System. After a 30 s incubation in >95% humidity, excess protein was blotted away for 2.5 s before being plunge frozen into liquid ethane using a Leica EM GP2 plunge freezer (Leica Microsystems). Frozen grids were imaged using a Titan Krios (Thermo Fisher) equipped with a K3 detector (Gatan). The cryoSPARC (*53*) software was used for data processing. Phenix (*54, 55*), Coot (*56*), Pymol (*57*), Chimera (*58*), ChimeraX (*59*) and Isolde (*60*) were used for model building and refinement.

### Mouse passive protection experiments

Twelve-month old female immunocompetent BALB/c mice purchased from Envigo (stock# 047) were used for passive immunization protection experiments. Mice were housed in groups of five animals per cage and fed standard chow diets. Mice were anesthetized with ketamine and xylazine prior to performing viral inoculations. Anesthetized mice were intranasally infected at 1X10^4^ PFU with RsSHC014-MA15 as described previously (*40*). For prophylaxis passive immunization experiments, mice were intraperitoneally injected with DH1338 at 15mg/kg or with flu IgG (CH65) at 15mg/kg 12 hours before infection. For the therapy passive immunization experiments, mice were intraperitoneally injected with DH1338 at 15mg/kg six hours post infection. Each treatment group contained N=9-10 mice. Following infection, mice were monitored daily for signs of disease. Ninety-six hours post infection, mice were sacrificed, lungs were harvested, and infectious virus replication was measured by plaque assays. All mouse studies were carried out in accordance with the recommendations for care and use of animals by the Office of Laboratory Animal Welfare (OLAW), National Institutes of Health and the Institutional Animal Care. Mouse studies performed at the University of North Carolina at Chapel Hill (Animal Welfare Assurance #A3410-01) using approved protocols by the Institutional Animal Care and Use Committee (IACUC). Mouse passive immunization and virus challenge experiments were performed in a biosafety level 3 (BSL3) facility at UNC Chapel Hill.

### Statistics Analysis

Data were plotted using Prism GraphPad 9.0. When appropriate, the area under the curve (AUC) was calculated using the linear trapezoidal method for longitudinal data (weeks 0-12). Wilcoxon rank-sum exact tests were performed to make comparisons between groups with an alpha level set at 0.05. The statistical analysis was performed using SAS 9.4 (SAS Institute). No adjustments were made to the alpha level for multiple comparisons, thus p < 0.05 is considered significant.

## Acknowledgments

We thank Margaret Deyton, Victoria Gee-Lai, Aja Sanzone, Nolan Jamieson, Lena Smith, Nicole De Naeyer and Conor Anderson for technical assistance. We thank Elizabeth Donahue, Cynthia Nagle and Kelly Soderberg for program management. We thank John Harrison, Alex Granados, Adrienne Goode, Anthony Cook, Alan Dodson, Katelyn Steingrebe, Bridget Bart, Laurent Pessaint, Alex VanRy, Daniel Valentin, Amanda Strasbaugh, and Mehtap Cabus for assistance with macaque studies. PCR purification for sequencing was performed by the DHVI Viral Genetics Analysis Core Facility. Single B cell sorting was performed in the DHVI Flow Cytometry Facility.

## Funding

This work was supported by funds from:

The State of North Carolina with funds from the federal CARES Act (to B.F.H.)

NIH, NIAID, DAIDS grant AI142596 (to B.F.H.)

NIH, NIAID, DAIDS grant AI158571 (to B.F.H.)

UC6-AI058607 (to G.D.S.)

NIH, NIAID, grants R01AI146101 and R01AI153064 (to N.P.)

The Ting Tsung & Wei Fong Chao Foundation (to B.F.H.)

Hanna H Gray Fellowship from the Howard Hughes Medical Institute (to D.R.M.)

Postdoctoral Enrichment Award from the Burroughs Wellcome Fund (to D.R.M.).

## Data and materials availability

All data are available in the main text or the supplementary materials. Electron microscopy data has been deposited under accession numbers in PDB: 8DPZ, 8DTK,7RAL. Materials are available with a Materials transfer agreement.

