## Supplemental figures for "Broadly neutralizing antibody induction by non-stabilized SARS-CoV-2 Spike mRNA vaccination in nonhuman primates"

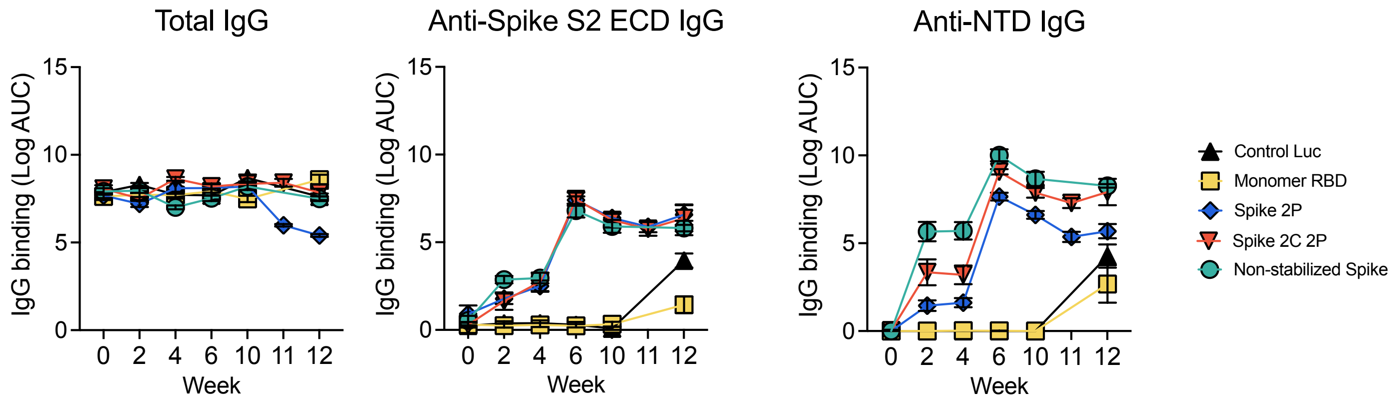

**Figure S1. Vaccine-induced SARS-CoV-2 specific IgG binding titers.** Total serum IgG binding magnitude to Spike 2P (S-2P), anti-Spike S2 ECD (ectodomain) and anti-NTD IgG were tested by ELISA and shown as log area under the curve (logAUC). Symbols indicate the group mean value ± SEM of three replicates.

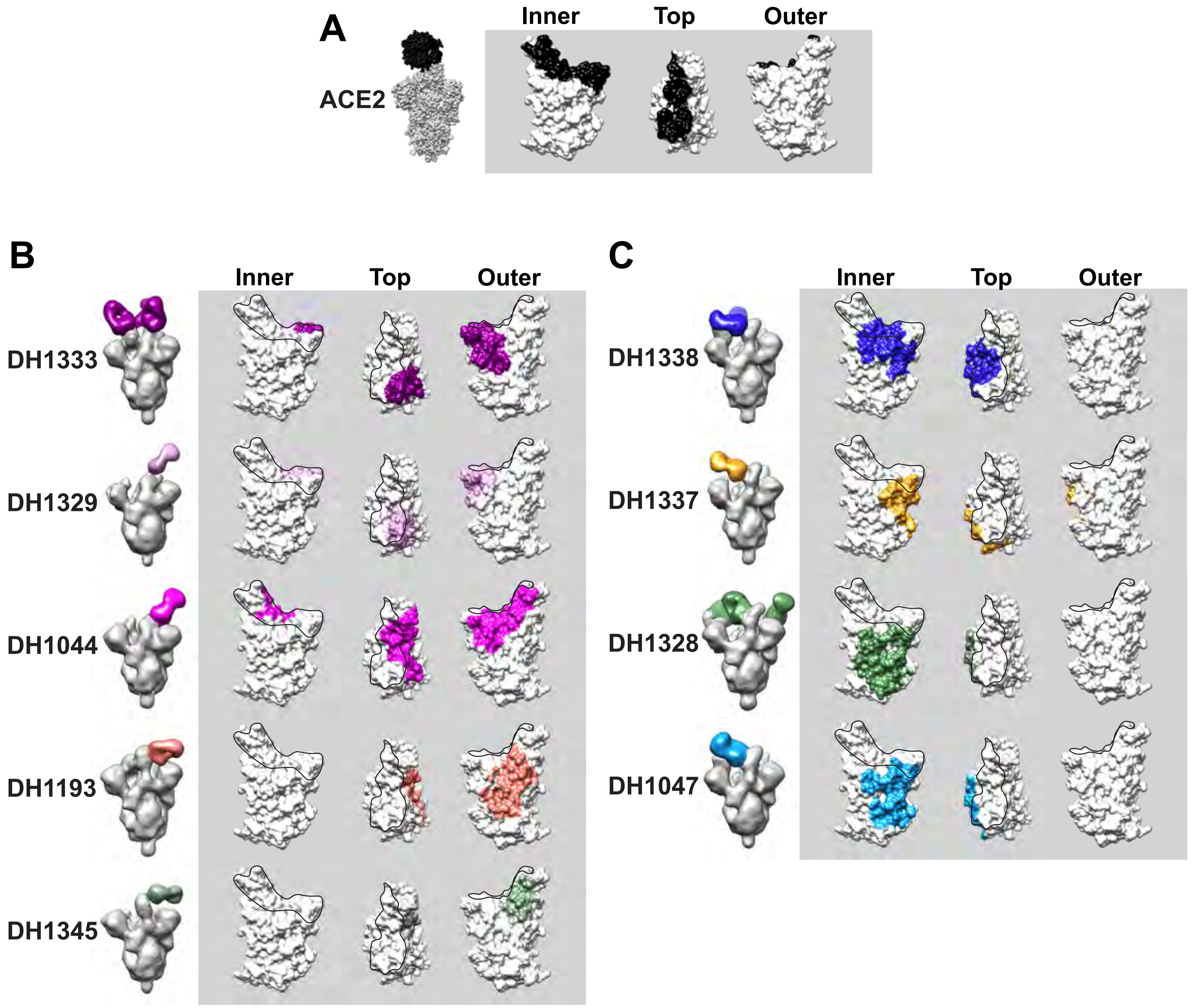

**Figure S2. Comparison of ACE2, macaque nAb, and human nAb binding footprints on SARS-CoV-2 RBD.** Antibody binding footprint was determined with 3D reconstruction of NSEM images of ACE2 or antibody bound to SARS-CoV-2 Spike. Spike is shown in gray with antibody Fab shown in different colors. The binding footprint on RBD for ACE2 or the specified antibody is shown to the right of the NSEM 3D reconstruction. (A) Structure of ACE2 (black) bound to SARS-CoV-2 Spike (gray). The ACE2 binding footprint is highlighted black on the RBD monomer to the right of the structure. (B,C) Binding footprint comparison for (B) outer and (C) inner domain antibodies. Human nAbs DH1044, DH1193, and DH1047 are shown for comparison to macaque antibodies (DH1333, DH1329, DH1345, DH1338, DH1337, and DH1328).

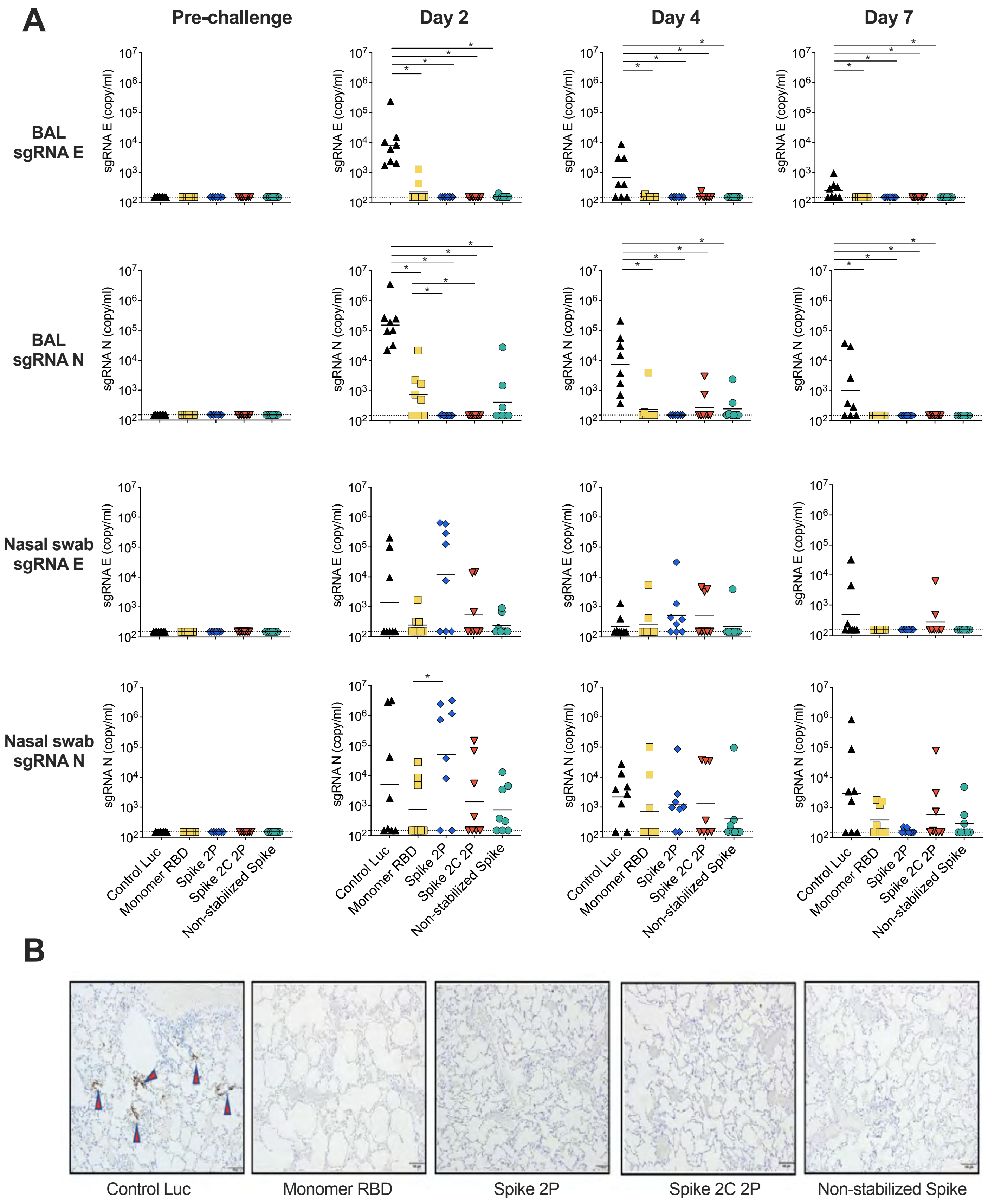
A
Pre-challenge
Day 2
Day 4
Day 7
BAL sgRNA E
BAL sgRNA N
Nasal swab sgRNA E
Nasal swab sgRNA N
sgRNA E (copy/ml)
sgRNA N (copy/ml)
Control Luc
Monomer RBD
Spike 2P
Spike 2C 2P
Non-stabilized Spike
B
Control Luc
Monomer RBD
Spike 2P
Spike 2C 2P
Non-stabilized Spike

**Figure S3. mRNA-LNP vaccination prevents virus replication in the lower respiratory tract after intranasal and intratracheal SARS-CoV-2 challenge in macaques.** (A) SARS-CoV-2 envelope gene (E gene) sgRNA and nucleocapsid gene (N gene) sgRNA in bronchoalveolar lavage (BAL) and nasal swab samples were quantitated pre-challenge, and on Day 2, 4, and 7. (B) A representative image of nucleocapsid antigen staining from each group of mRNA-LNP vaccinated macaques is shown. All images are shown at 10x magnification. Scale bars, 100 μm.

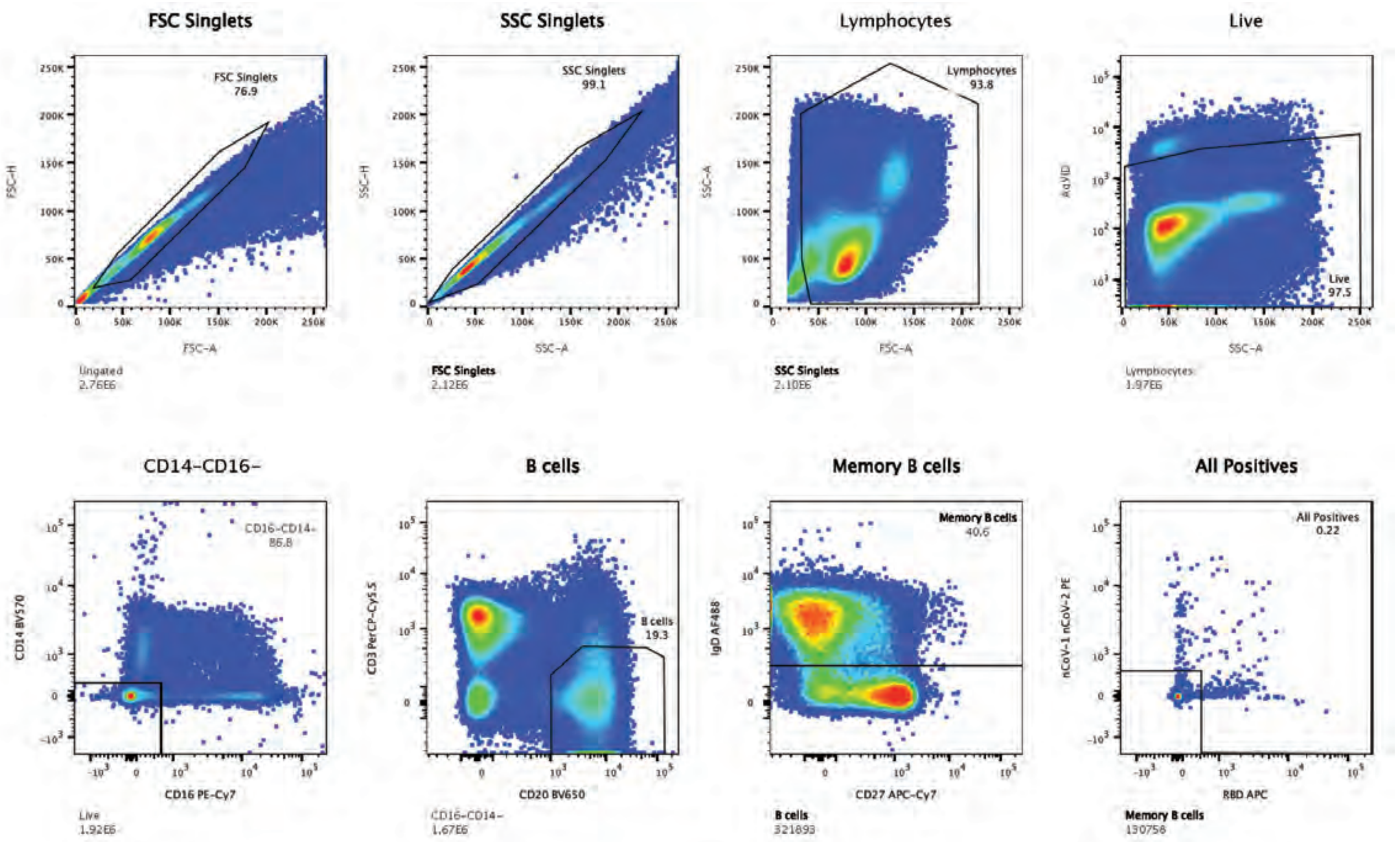

**Figure S4. Flow cytometric gating strategies for antigen-specific B cell isolation.** The gating strategy to identify SARS-CoV-2 full length spike and RBD-specific memory B cells.

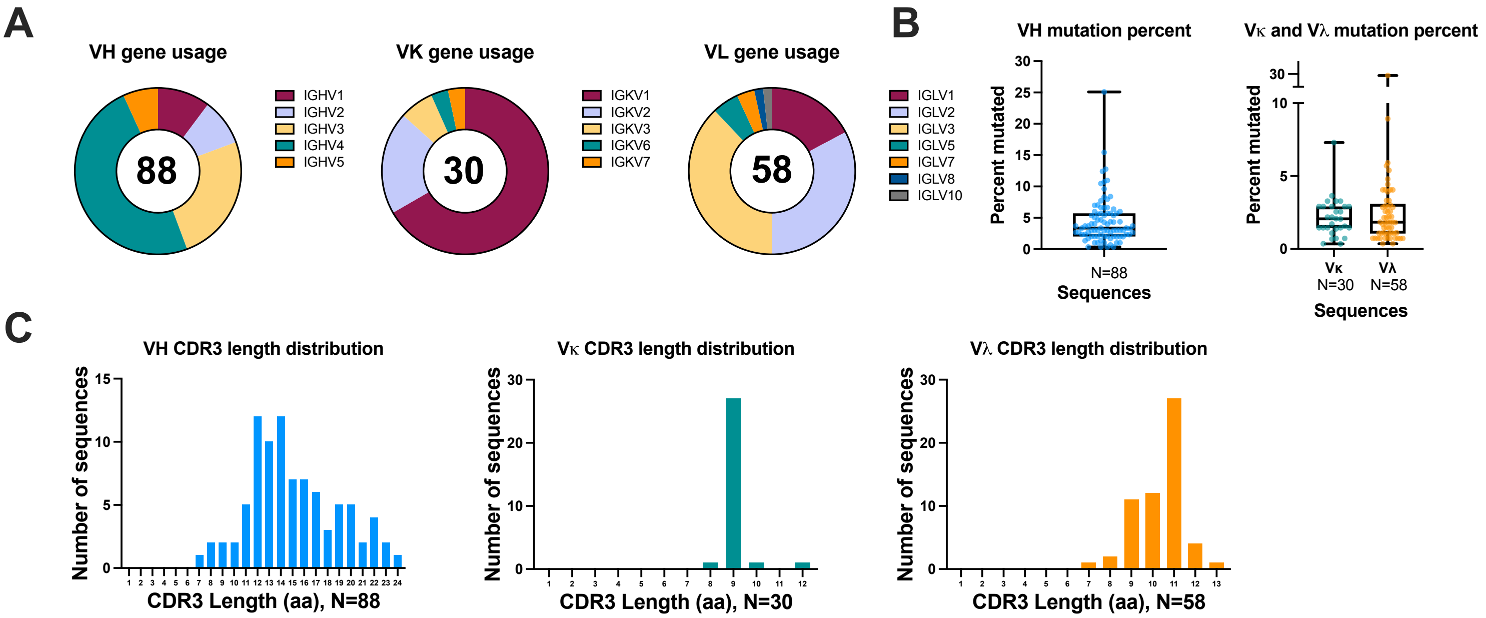

D

| Antibody ID | VH gene | VH CDR3 length | VH mutation % | VL gene | VL CDR3 length | VL mutation % |
|---|---|---|---|---|---|---|
| DH1323 | IGHV1-c*01 | 16 | 3.020 | IGKV1-b*06 | 9 | 1.460 |
| DH1334 | IGHV3-x*04 | 17 | 12.752 | IGLV3-j*01 | 10 | 0.738 |
| DH1340 | IGHV4-j*02 | 16 | 2.685 | IGLV2-j*15 | 7 | 0.738 |
| DH1341 | IGHV4-m*02 | 13 | 6.579 | IGLV1-i*02 | 11 | 2.166 |
| DH1330 | IGHV2-a*02 | 16 | 1.329 | IGKV1-g*04 | 9 | 0.730 |
| DH1324 | IGHV4-j*02 | 17 | 2.349 | IGKV1-r*01 | 12 | 1.460 |
| DH1338 | IGHV4-j*02 | 21 | 1.678 | IGKV3-b*19 | 9 | 3.650 |
| DH1335 | IGHV4-b*01 | 15 | 4.362 | IGLV3-j*05 | 11 | 1.107 |
| DH1333 | IGHV3-m*05 | 15 | 2.961 | IGLV7-c*01 | 10 | 0.714 |
| DH1325 | IGHV4-n*01 | 14 | 2.658 | IGLV2-j*15 | 11 | 3.321 |
| DH1329 | IGHV1-c*01 | 16 | 4.027 | IGKV3-d*05 | 9 | 1.460 |
| DH1328 | IGHV4-n*03 | 23 | 1.993 | IGKV1-g*04 | 9 | 0.365 |
| DH1345 | IGHV5-a*04 | 12 | 2.349 | IGLV1-a*03 | 11 | 1.805 |
| DH1342 | IGHV4-n*03 | 14 | 6.645 | IGLV3-e*01 | 11 | 1.107 |
| DH1336 | IGHV4-e*01 | 14 | 7.718 | IGKV2-g*01 | 9 | 2.076 |
| DH1326 | IGHV2-b*04 | 22 | 1.993 | IGLV2-j*16 | 12 | 4.059 |
| DH1337 | IGHV4-e*01 | 20 | 5.705 | IGLV1-i*02 | 12 | 1.083 |
| DH1339 | IGHV4-j*02 | 14 | 3.356 | IGLV2-d*07 | 10 | 8.929 |
| DH1331 | IGHV3-j*02 | 14 | 2.961 | IGLV2-j*15 | 11 | 2.214 |
| DH1343 | IGHV5-a*03 | 16 | 1.007 | IGLV1-a*01 | 11 | 1.444 |
| DH1332 | IGHV3-j*02 | 13 | 0.987 | IGLV3-j*05 | 11 | 3.321 |
| DH1327 | IGHV3-k*04 | 10 | 2.303 | IGKV3-b*19 | 8 | 2.190 |
| DH1344 | IGHV5-a*04 | 11 | 2.013 | IGLV1-a*01 | 12 | 1.083 |

**Figure S5. Monoclonal antibody isolation from wildtype SARS-CoV-2 spike mRNA-LNP vaccinated rhesus macaques.** (A) Frequencies of VH, VK, VL gene segment usage of 88 non-clonal sequences of isolated spike protein polyreactive antibodies from mRNA-LNP immunized macaques. (B) Frequencies of VH, VK, VL CDRH3 length of 88 non-clonal sequences of isolated spike protein reactive antibodies from mRNA-LNP immunized macaques. (C) Distribution of mutation rates for heavy and light chains of 88 non-clonal sequences of isolated spike protein reactive antibodies from mRNA-LNP immunized macaques. (D) Immunogenetic analysis of the monoclonal antibodies isolated from wild-type spike mRNA-LNP vaccination in rhesus macaques.

| | Antigens | | | | | | | | | | | | | | | | | | |
|---|---|---|---|---|---|---|---|---|---|---|---|---|---|---|---|---|---|---|---|
| Antibody ID | SARS-CoV-2 (2019-nCoV) Spike Protein (S1+S2 ECD) | SARS-CoV-2 Spike protein (RBD) | SARS-CoV-2 (2019-nCoV) Spike Protein (S2 ECD) | SARS-CoV Spike Protein DeltaTM Recombinant | SARS-CoV, WH20 Coronavirus spike S1 | SARS-CoV-1 RBD | MERS-CoV Coronavirus spike RBD | MERS-CoV Coronavirus spike S2 | MERS-CoV, Coronavirus spike S1+S2 | Recombinant MERS-CoV Spike/S1 Protein (S1 subunit) | HCoV-NL63 Spike Protein (S1+S2 ECD) | HCoV-229E Spike Protein (S1+S2 ECD) | Human Coronavirus HKU1 (isolate N5) (HCoV-HKU1) Spike Protein (S1+S2 ECD) | HCoV-OC43 Spike Protein (S1+S2 ECD) | nCOV-1 nCOV-2p | NTD | Streptavidin control | biotin-Man9 V3 (negative control) | Candida Albicans |
| DH1323 | 0.204 | 0.902 | 0.003 | 0.031 | -0.020 | -0.003 | 0.195 | -0.031 | -0.004 | -0.008 | -0.020 | -0.017 | -0.009 | -0.012 | -0.029 | 0.005 | 0.000 | 0.021 | 0.011 |
| DH1324 | 0.009 | 0.109 | 0.023 | 0.001 | -0.014 | 0.015 | -0.027 | -0.028 | -0.008 | -0.009 | -0.033 | -0.025 | -0.022 | -0.032 | -0.046 | -0.009 | 0.000 | -0.032 | 0.073 |
| DH1325 | 2.611 | 3.303 | 0.032 | 0.075 | 0.726 | 0.188 | -0.014 | -0.018 | 0.006 | 0.003 | -0.008 | -0.024 | -0.012 | -0.011 | 0.257 | 0.002 | 0.000 | -0.020 | 0.052 |
| DH1326 | 2.930 | 3.327 | 0.022 | 0.533 | 2.780 | 3.229 | 0.004 | 0.002 | 0.019 | 0.009 | -0.001 | -0.006 | 0.025 | 0.012 | 0.423 | 0.168 | 0.000 | 0.005 | 0.047 |
| DH1327 | 3.013 | 3.260 | 0.039 | -0.004 | 0.011 | 0.007 | -0.021 | -0.018 | -0.003 | -0.009 | -0.018 | -0.024 | -0.006 | -0.012 | 0.336 | 0.002 | 0.000 | -0.004 | 0.009 |
| DH1328 | -0.007 | 0.008 | -0.010 | -0.008 | -0.004 | 0.019 | -0.017 | -0.019 | 0.004 | 0.031 | -0.030 | -0.026 | -0.007 | -0.004 | -0.072 | -0.053 | 0.000 | -0.012 | 0.007 |
| DH1329 | 3.287 | 3.507 | 0.003 | 0.126 | 0.558 | 0.721 | -0.003 | 0.007 | 0.007 | 0.012 | 0.029 | 0.009 | 0.031 | 0.029 | 1.059 | 0.036 | 0.000 | -0.049 | 0.003 |
| DH1330 | 3.089 | 3.502 | 0.035 | 0.025 | 0.092 | 0.291 | -0.021 | -0.007 | -0.006 | 0.005 | 0.013 | -0.005 | -0.009 | -0.002 | 0.705 | 0.095 | 0.000 | -0.045 | -0.015 |
| DH1331 | 2.455 | 3.384 | -0.006 | 0.000 | 0.257 | 2.085 | -0.002 | -0.004 | -0.003 | 0.010 | 0.011 | 0.012 | 0.007 | 0.006 | 0.369 | 0.058 | 0.000 | -0.026 | 0.033 |
| DH1332 | 2.820 | 3.446 | 0.279 | -0.016 | 0.005 | 0.067 | 0.168 | 0.040 | -0.002 | 0.001 | 0.002 | -0.014 | 0.035 | -0.004 | 0.300 | 0.087 | 0.000 | -0.036 | 0.024 |
| DH1333 | 2.945 | 3.410 | 0.085 | 0.077 | 0.144 | 0.144 | 0.080 | 0.133 | 0.082 | 0.080 | 0.094 | 0.127 | 0.126 | 0.120 | 0.873 | 0.011 | 0.000 | -0.142 | -0.355 |
| DH1334 | 2.077 | 3.080 | 0.004 | 0.283 | 2.204 | 2.448 | -0.011 | -0.004 | 0.002 | 0.024 | 0.021 | -0.014 | -0.014 | -0.003 | 0.500 | 1.856 | 0.000 | -0.040 | 0.002 |
| DH1335 | 2.990 | 3.449 | 0.105 | 1.181 | 3.188 | 3.360 | -0.002 | 0.039 | 0.041 | 0.006 | 0.017 | 0.019 | 0.030 | 0.031 | 0.465 | 1.005 | 0.000 | -0.043 | 0.046 |
| DH1336 | 0.305 | 0.967 | 0.032 | -0.012 | -0.014 | 0.038 | -0.006 | 0.002 | 0.005 | -0.001 | -0.003 | -0.006 | -0.006 | -0.016 | 0.087 | 0.073 | 0.000 | -0.041 | -0.016 |
| DH1337 | 2.591 | 3.228 | 0.043 | 1.055 | 2.941 | 3.190 | -0.012 | 0.030 | 0.024 | 0.010 | 0.027 | 0.035 | 0.065 | 0.079 | 0.284 | 0.298 | 0.000 | -0.128 | -0.083 |
| DH1338 | 2.846 | 3.451 | 0.019 | 1.981 | 3.346 | 3.495 | 0.025 | 0.022 | 0.023 | 0.029 | 0.067 | 0.027 | 0.043 | 0.031 | 0.624 | 0.012 | 0.000 | -0.014 | 0.534 |
| DH1339 | 0.002 | 0.088 | 0.052 | 0.005 | -0.011 | 0.039 | 0.009 | 0.009 | 0.049 | 0.012 | 0.010 | 0.010 | 0.025 | 0.024 | -0.013 | 0.109 | 0.000 | -0.045 | 0.141 |
| DH1340 | 2.981 | 3.196 | 0.053 | 0.017 | 0.120 | 0.133 | 0.049 | 0.022 | 0.044 | 0.043 | 0.049 | 0.028 | 0.052 | 0.055 | 0.588 | 0.340 | 0.000 | -0.078 | 0.353 |
| DH1341 | 3.028 | 3.378 | -0.003 | 0.170 | 1.678 | 1.598 | 0.006 | 0.003 | 0.020 | 0.013 | 0.026 | 0.023 | 0.025 | 0.025 | 0.689 | 0.019 | 0.000 | -0.069 | 0.132 |
| DH1342 | 1.539 | 3.233 | 0.060 | 0.008 | 0.038 | 0.035 | 0.034 | 0.050 | 0.019 | 0.028 | 0.032 | 0.044 | 0.045 | 0.019 | 0.196 | 0.080 | 0.000 | -0.031 | 0.027 |
| DH1343 | 2.610 | 3.301 | 0.006 | -0.005 | -0.028 | 0.028 | 0.021 | 0.010 | 0.032 | 0.046 | 0.036 | 0.009 | 0.041 | 0.060 | 0.479 | 0.595 | 0.000 | 0.060 | 0.275 |
| DH1344 | 2.345 | 3.344 | 0.043 | 0.033 | 0.005 | 0.063 | 0.050 | 0.045 | 0.052 | 0.117 | 0.181 | 0.089 | 0.062 | 0.036 | 0.960 | -0.026 | 0.000 | 0.023 | -0.081 |
| DH1345 | 3.155 | 3.490 | 3.144 | 0.040 | 0.152 | 0.263 | 0.040 | 0.057 | 0.133 | 0.028 | 0.028 | 0.044 | 0.054 | 0.036 | 0.543 | 0.497 | 0.000 | -0.063 | 0.212 |
| DH1453 | 3.125 | 0.032 | 0.005 | 0.924 | 3.105 | 0.011 | -0.011 | -0.002 | -0.014 | 0.004 | 0.002 | 0.003 | 0.009 | 0.009 | 0.815 | 3.366 | 0.000 | -0.037 | -0.013 |
| DH1454 | 1.297 | 0.047 | -0.008 | 0.064 | 1.141 | 0.037 | -0.033 | 0.004 | 0.029 | -0.006 | 0.022 | -0.002 | -0.022 | -0.016 | 0.084 | 3.216 | 0.000 | -0.009 | 0.080 |
| DH1455 | 0.034 | -0.004 | -0.012 | -0.001 | -0.036 | 0.004 | -0.011 | 0.007 | -0.002 | 0.009 | 0.034 | 0.013 | 0.012 | 0.017 | -0.022 | 0.359 | 0.000 | -0.033 | -0.011 |
| DH1456 | 1.972 | 0.355 | 0.017 | 0.004 | 0.020 | 0.052 | 0.004 | 0.009 | 0.025 | 0.017 | 0.021 | 0.013 | 0.025 | 0.021 | 0.289 | 3.077 | 0.000 | -0.066 | 0.011 |
| DH1457 | 0.124 | 0.134 | 0.007 | -0.018 | -0.029 | 0.002 | -0.012 | 0.001 | 0.001 | 0.000 | -0.003 | 0.000 | 0.005 | 0.038 | 0.035 | 0.772 | 0.000 | -0.174 | -0.150 |
| DH1458 | 1.476 | 0.070 | -0.013 | 0.051 | 0.035 | 0.147 | -0.006 | 0.026 | 0.031 | 0.006 | 0.003 | 0.004 | 0.016 | 0.017 | 0.689 | 3.289 | 0.000 | -0.007 | -0.024 |
| DH1459 | 0.188 | 0.275 | 0.019 | 0.044 | 0.243 | 0.278 | -0.012 | 0.010 | 0.004 | 0.003 | 0.007 | 0.004 | -0.001 | 0.022 | -0.026 | 0.009 | 0.000 | -0.036 | -0.001 |
| DH1460 | 0.009 | 0.100 | -0.023 | -0.017 | -0.058 | 0.023 | 0.012 | 0.025 | 0.013 | 0.002 | 0.004 | 0.009 | -0.016 | -0.002 | 0.123 | 2.720 | 0.000 | -0.262 | -0.244 |
| DH1461 | -0.003 | 0.037 | -0.032 | -0.019 | -0.058 | 0.003 | 0.010 | -0.011 | -0.017 | 0.000 | -0.006 | -0.009 | -0.019 | 0.000 | 0.428 | 3.298 | 0.000 | -0.020 | 0.012 |
| DH1462 | 1.962 | 0.046 | 0.361 | 0.000 | 0.019 | 0.052 | 0.020 | 0.022 | 0.023 | 0.011 | 0.012 | 0.014 | 0.027 | 0.035 | 0.546 | 3.282 | 0.000 | -0.018 | 0.023 |
| DH1463 | 0.530 | 1.658 | -0.011 | -0.020 | 0.097 | 0.146 | -0.020 | 0.000 | 0.002 | 0.005 | -0.009 | -0.004 | -0.023 | -0.015 | 0.973 | 3.299 | 0.000 | -0.035 | -0.013 |
| DH1464 | 0.785 | 0.023 | 0.033 | 0.005 | -0.022 | 0.021 | -0.007 | -0.005 | 0.001 | 0.009 | 0.003 | 0.004 | 0.010 | 0.006 | 0.557 | 3.111 | 0.000 | 0.001 | 0.106 |
| DH1465 | 0.066 | 0.052 | 0.010 | -0.012 | -0.011 | 0.009 | 0.001 | 0.041 | 0.042 | 0.010 | 0.000 | 0.029 | 0.025 | 0.031 | 0.487 | 3.211 | 0.000 | -0.027 | -0.004 |
| DH1466 | 0.029 | 0.002 | -0.014 | -0.004 | 0.009 | 0.004 | -0.027 | -0.007 | -0.005 | 0.002 | -0.002 | -0.003 | -0.019 | -0.015 | 0.337 | 1.887 | 0.000 | -0.254 | -0.252 |
| DH1467 | 0.015 | 0.014 | -0.009 | 0.005 | -0.001 | 0.012 | -0.006 | -0.019 | -0.014 | 0.002 | 0.010 | 0.031 | -0.011 | 0.025 | -0.041 | 0.334 | 0.000 | 0.010 | 0.033 |
| DH1468 | 2.862 | 3.584 | 0.106 | 0.091 | 0.071 | 0.145 | 0.097 | 0.075 | 0.117 | 0.099 | 0.066 | 0.104 | 0.107 | 0.059 | 0.538 | -0.069 | 0.000 | -0.206 | 0.100 |
| DH1469 | 2.929 | 3.431 | 0.005 | 0.013 | -0.021 | 0.043 | -0.011 | 0.008 | 0.005 | 0.001 | -0.003 | 0.000 | -0.014 | -0.014 | 0.530 | 0.026 | 0.000 | -0.038 | 0.007 |
| DH1470 | 2.718 | 3.305 | -0.024 | -0.018 | 0.022 | 0.003 | -0.023 | -0.008 | -0.024 | 0.004 | -0.004 | -0.012 | -0.018 | -0.018 | 0.511 | 0.727 | 0.000 | -0.056 | -0.026 |
| DH1471 | 1.209 | 2.813 | 0.312 | 0.055 | 0.128 | 0.067 | 0.105 | 0.076 | 0.080 | 0.073 | 0.048 | 0.164 | 0.288 | 0.081 | -0.015 | 0.159 | 0.000 | 0.005 | 0.316 |
| DH1472 | 1.810 | 3.171 | -0.005 | -0.003 | 0.009 | 0.011 | -0.018 | -0.018 | -0.003 | -0.002 | -0.024 | -0.014 | -0.011 | -0.003 | 0.120 | -0.001 | 0.000 | -0.001 | -0.002 |
| DH1473 | 0.128 | 0.159 | -0.003 | 0.015 | 0.108 | 0.176 | -0.002 | 0.043 | 0.024 | 0.039 | 0.000 | 0.029 | 0.041 | 0.049 | 0.133 | 0.068 | 0.000 | -0.024 | 0.009 |
| DH1474 | 1.070 | 2.250 | 0.043 | 0.007 | 0.005 | 0.058 | 0.038 | 0.052 | 0.043 | 0.022 | 0.020 | 0.025 | 0.086 | 0.054 | 0.206 | 0.245 | 0.000 | -0.093 | -0.081 |
| DH1475 | 2.304 | 3.481 | 0.009 | 1.238 | 3.268 | 3.442 | -0.014 | -0.027 | -0.006 | -0.004 | -0.014 | 0.054 | -0.017 | -0.017 | 0.117 | 0.003 | 0.000 | -0.015 | 0.015 |
| DH1476 | 2.878 | 3.335 | -0.010 | 1.509 | 3.134 | -0.001 | -0.014 | -0.008 | 0.000 | 0.007 | -0.023 | -0.021 | 0.356 | -0.004 | 0.198 | 0.007 | 0.000 | 0.052 | 0.044 |
| DH1477 | 0.629 | 0.040 | 0.746 | 0.089 | -0.006 | 0.021 | 0.025 | 0.027 | 0.026 | 0.030 | 0.015 | 0.063 | 0.041 | 0.026 | 0.487 | -0.005 | 0.000 | -0.053 | 0.160 |
| DH1478 | 3.228 | 0.755 | 3.374 | 0.169 | 0.230 | 0.235 | 0.008 | 0.041 | 0.171 | 0.063 | 0.048 | 0.075 | 0.109 | 0.106 | 0.178 | 0.256 | 0.000 | -0.212 | 0.768 |
| DH1479 | -0.023 | 0.007 | -0.024 | -0.017 | -0.058 | 0.004 | 0.004 | 0.013 | 0.003 | 0.001 | -0.011 | -0.012 | -0.008 | -0.004 | 0.011 | 2.103 | 0.000 | -0.024 | 0.004 |
| DH1480 | 0.372 | 0.370 | 0.329 | 0.220 | 0.304 | 0.346 | 0.413 | 0.307 | 0.265 | 0.245 | 0.246 | 0.269 | 0.340 | 0.344 | 0.037 | 1.689 | 0.000 | 0.228 | 0.138 |
| DH1481 | 0.805 | 0.093 | 0.048 | 0.068 | 0.118 | 0.113 | 0.051 | 0.031 | 0.088 | 0.044 | 0.049 | 0.084 | 0.091 | 0.056 | 0.076 | 1.939 | 0.000 | 0.014 | 0.104 |
| DH1482 | 2.712 | 3.302 | -0.002 | 0.400 | 2.358 | 0.008 | -0.019 | 0.001 | 0.001 | 0.024 | 0.010 | 0.028 | 0.011 | 0.031 | 1.034 | 3.721 | 0.000 | -0.016 | 0.126 |
| DH1483 | 2.056 | 3.171 | -0.011 | 0.039 | -0.016 | -0.002 | -0.008 | -0.017 | -0.005 | -0.001 | -0.029 | -0.032 | -0.012 | -0.008 | 0.118 | 0.015 | 0.000 | -0.006 | 0.017 |
| DH1484 | 2.459 | 3.100 | 0.004 | 0.005 | -0.057 | 0.034 | -0.021 | 0.013 | 0.005 | 0.000 | -0.006 | -0.009 | 0.007 | 0.006 | 0.388 | 0.042 | 0.000 | -0.057 | 0.025 |
| DH1485 | 2.519 | 3.158 | 0.019 | 0.002 | 0.000 | 0.044 | -0.023 | -0.027 | -0.010 | 0.010 | -0.027 | -0.032 | 0.039 | 0.018 | 0.157 | 0.045 | 0.000 | 0.008 | 0.025 |
| DH1486 | -0.020 | 0.000 | -0.003 | 0.019 | 0.040 | 0.015 | -0.011 | 0.033 | -0.005 | -0.004 | -0.008 | -0.006 | -0.008 | -0.012 | -0.025 | 0.102 | 0.000 | -0.032 | 0.024 |
| DH1487 | 0.125 | 0.064 | 0.010 | 0.002 | -0.014 | 0.033 | 0.015 | 0.019 | 0.006 | 0.000 | -0.004 | -0.004 | 0.004 | 0.004 | 0.056 | 2.179 | 0.000 | -0.041 | 0.000 |

**Figure S6. High-throughput ELISA binding screen of antibodies recovered from wildtype SARS-CoV-2 spike mRNA-LNP vaccinated rhesus macaques.** Antibodies were tested in unpurified cell culture supernatant at a single dilution. Yellow highlighted values are above 0.1 OD450nm and are considered positive.

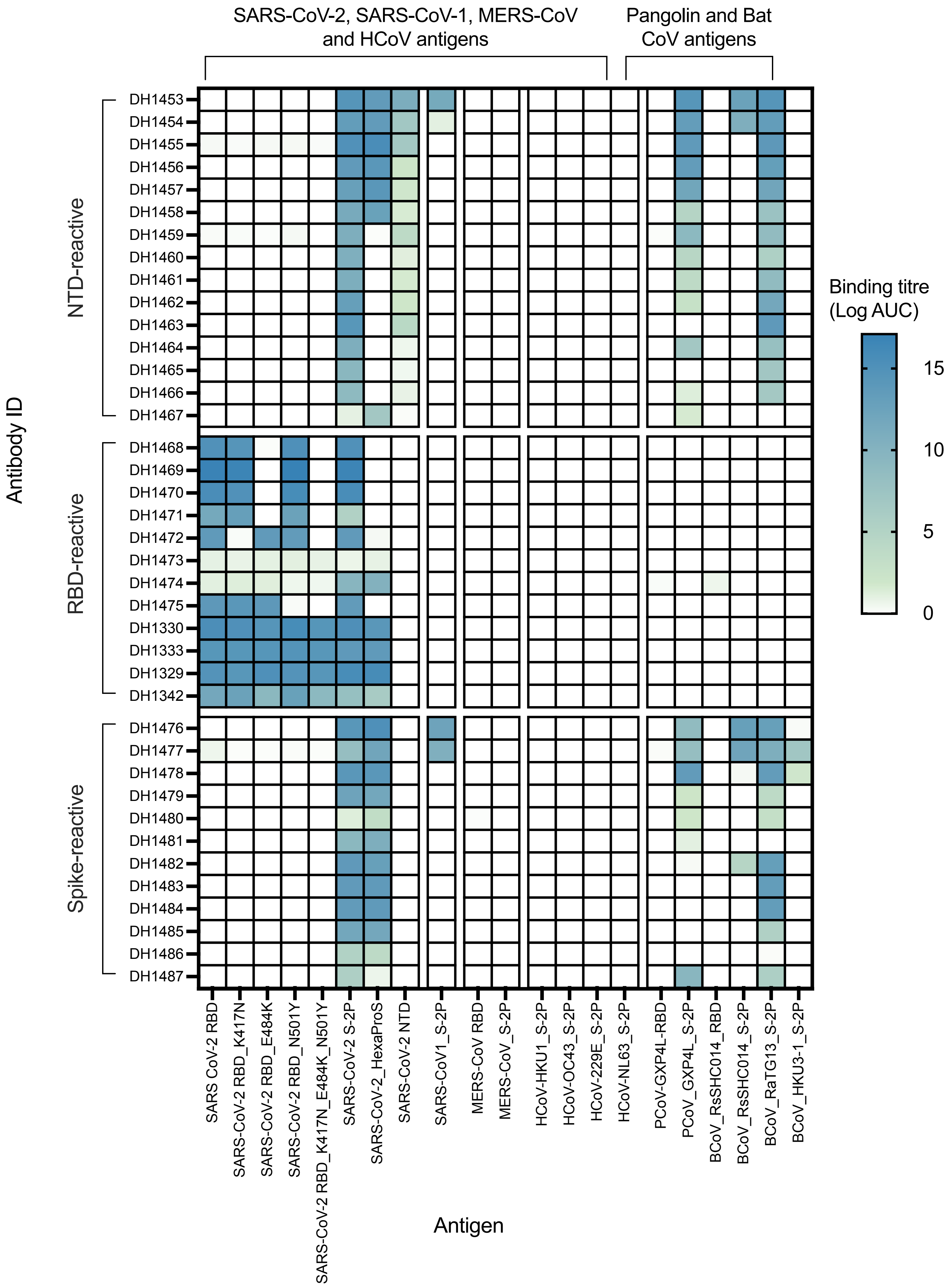

SARS-CoV-2, SARS-CoV-1, MERS-CoV and HCoV antigens
Pangolin and Bat CoV antigens
Antibody ID
NTD-reactive
RBD-reactive
Spike-reactive
DH1453
DH1454
DH1455
DH1456
DH1457
DH1458
DH1459
DH1460
DH1461
DH1462
DH1463
DH1464
DH1465
DH1466
DH1467
DH1468
DH1469
DH1470
DH1471
DH1472
DH1473
DH1474
DH1475
DH1330
DH1333
DH1329
DH1342
DH1476
DH1477
DH1478
DH1479
DH1480
DH1481
DH1482
DH1483
DH1484
DH1485
DH1486
DH1487
SARS CoV-2 RBD
SARS-CoV-2 RBD_K417N
SARS-CoV-2 RBD_E484K
SARS-CoV-2 RBD_N501Y
SARS-CoV-2 RBD_K417N_E484K_N501Y
SARS-CoV-2 S-2P
SARS-CoV-2_HexaProS
SARS-CoV-2 NTD
SARS-CoV1_S-2P
MERS-CoV RBD
MERS-CoV_S-2P
HCoV-HKU1_S-2P
HCoV-OC43_S-2P
HCoV-229E_S-2P
HCoV-NL63_S-2P
PCoV-GXP4L-RBD
PCoV_GXP4L_S-2P
BCoV_RsSHC014_RBD
BCoV_RsSHC014_S-2P
BCoV_RaTG13_S-2P
BCoV_HKU3-1_S-2P
Antigen
Binding titre (Log AUC)
15
10
5
0

**Figure S7. Monoclonal antibodies isolated from wildtype S-tm mRNA-LNP immunized rhesus macaques bind SARS-CoV-2 variants and SARS-related betacoronaviruses and possess different epitope specificities.** Antibodies labeled Spike-reactive bound to various domains of Spike in initial binding screens and confirmatory ELISA.

# A

Competitive relationship between vaccine-induced rhesus macaque mAbs with vaccine or infection induced human/ mouse mAbs

| Group | mAb | DH1041 | DH1042 | DH1044 | DH1047 | DH1073 | DH1235 | DH1284 | DH1193 | S309 | CR3022 | SP1-77 |
|---|---|---|---|---|---|---|---|---|---|---|---|---|
| Outer RBD face macaque antibodies | DH1333 | 1.322 | 1.223 | 0.508 | 10.000 | 10.000 | 10.000 | 10.000 | 0.329 | 10.000 | 10.000 | 1.014 |
| | DH1345 | 10.000 | 10.000 | 0.425 | 10.000 | 8.864 | 10.000 | 10.000 | 0.208 | 10.000 | 10.000 | 10.000 |
| | DH1341 | 10.000 | 0.605 | 0.309 | 10.000 | 1.223 | 10.000 | 10.000 | 0.231 | 0.985 | 10.000 | 10.000 |
| | DH1344 | 10.000 | 10.000 | 0.562 | 10.000 | 10.000 | 10.000 | 10.000 | 0.248 | 10.000 | 10.000 | 10.000 |
| | DH1343 | 10.000 | 10.000 | 0.685 | 10.000 | 10.000 | 10.000 | 10.000 | 0.326 | 10.000 | 10.000 | 10.000 |
| | DH1329 | 0.417 | 0.421 | 0.214 | 10.000 | 10.000 | 10.000 | 10.000 | 10.000 | 0.522 | 10.000 | 1.021 |
| | DH1330 | 0.944 | 0.705 | 0.334 | 10.000 | 10.000 | 10.000 | 10.000 | 0.214 | 10.000 | 10.000 | 10.000 |
| | DH1325 | 10.000 | 10.000 | 0.501 | 10.000 | 10.000 | 10.000 | 10.000 | 0.287 | 10.000 | 10.000 | 10.000 |
| | DH1340 | 10.000 | 10.000 | 0.569 | 10.000 | 4.369 | 10.000 | 10.000 | 0.300 | 10.000 | 10.000 | 1.200 |
| | DH1335 | 10.000 | 10.000 | 0.537 | 10.000 | 10.000 | 10.000 | 10.000 | 0.224 | 10.000 | 10.000 | 10.000 |
| Inner RBD face macaque antibodies | DH1338 | 10.000 | 10.000 | 10.000 | 0.346 | 10.000 | 0.215 | 10.000 | 10.000 | 10.000 | 0.086 | 10.000 |
| | DH1328 | 10.000 | 10.000 | 10.000 | 0.384 | 10.000 | 0.217 | 10.000 | 10.000 | 10.000 | 0.104 | 10.000 |
| | DH1337 | 10.000 | 10.000 | 10.000 | 0.782 | 10.000 | 0.676 | 10.000 | 10.000 | 10.000 | 0.236 | 10.000 |
| | DH1324 | 10.000 | 10.000 | 10.000 | 0.539 | 10.000 | 0.475 | 10.000 | 10.000 | 10.000 | 0.237 | 10.000 |
| | DH1336 | 10.000 | 10.000 | 10.000 | 1.380 | 10.000 | 10.000 | 10.000 | 10.000 | 10.000 | 0.171 | 10.000 |
| | DH1326 | 10.000 | 10.000 | 10.000 | 1.087 | 10.000 | 0.528 | 10.000 | 10.000 | 10.000 | 0.183 | 10.000 |
| | DH1339 | 10.000 | 10.000 | 10.000 | 4.504 | 10.000 | 3.078 | 10.000 | 10.000 | 10.000 | 0.981 | 10.000 |
| | DH1331 | 10.000 | 10.000 | 10.000 | 10.000 | 10.000 | 10.000 | 10.000 | 10.000 | 10.000 | 0.260 | 10.000 |
| | DH1327 | 10.000 | 10.000 | 10.000 | 10.000 | 10.000 | 10.000 | 10.000 | 10.000 | 10.000 | 2.244 | 10.000 |
| | DH1332 | 10.000 | 10.000 | 10.000 | 10.000 | 10.000 | 10.000 | 10.000 | 10.000 | 10.000 | 10.000 | 10.000 |
| | DH1334 | 10.000 | 10.000 | 10.000 | 10.000 | 10.000 | 10.000 | 10.000 | 10.000 | 10.000 | 10.000 | 10.000 |
| | DH1342 | 10.000 | 10.000 | 10.000 | 10.000 | 10.000 | 10.000 | 10.000 | 10.000 | 10.000 | 10.000 | 10.000 |

$IC_{50}$ (ug/mL): <0.1 | <1 | <10 | <5 | >10

# B

| mAb | Live virus neutralization | | | Pseudovirus neutraliazation (FDA) | | | | | | | | | Psedovirus neutraliazation (Duke) | |
|---|---|---|---|---|---|---|---|---|---|---|---|---|---|---|
| | SARS-CoV-2 D614G | SARS-CoV | RsSHC014 | WA-1 | D614G | B.1.1.7 (Alpha) | B.1.351 (Beta) | P.1 (Gamma) | B.1.617.2 (Delta) | B.1.617.1 (Kappa) | B.1.429 (Epsilon) | B.1.526 (Iota) | BA.1 (Omicron) | BA.4/BA.5 |
| DH1333 | 0.006 | 10.000 | 10.000 | 0.016 | 0.010 | 0.030 | 0.022 | 0.028 | 0.025 | 0.016 | 0.016 | 0.020 | 50.000 | 0.030 |
| DH1345 | 1.231 | 10.000 | 10.000 | 0.141 | 0.660 | 0.157 | 0.611 | 0.177 | 0.315 | 0.326 | 0.156 | 0.515 | 1.600 | 3.400 |
| DH1341 | 6.368 | 10.000 | 10.000 | 1.401 | 3.600 | 1.481 | 3.300 | 1.658 | 2.924 | 1.647 | 1.304 | 2.755 | 3.300 | 12.000 |
| DH1344 | 0.220 | 10.000 | 10.000 | 0.444 | 3.900 | 0.572 | 1.374 | 0.654 | 0.951 | 1.414 | 0.560 | 1.112 | 28.000 | 21.000 |
| DH1343 | 1.107 | 10.000 | 10.000 | 0.533 | 2.500 | 0.588 | 1.255 | 0.589 | 0.913 | 1.127 | 0.621 | 1.443 | 18.000 | 25.000 |
| DH1329 | 0.003 | 10.000 | 10.000 | 0.023 | 0.010 | 0.023 | 0.022 | 0.024 | 10.000 | 10.000 | 10.000 | 0.027 | 0.280 | 50.000 |
| DH1330 | 0.206 | 10.000 | 10.000 | 0.120 | 0.330 | 0.143 | 2.793 | 0.538 | 10.000 | 10.000 | 10.000 | 0.904 | 4.300 | 50.000 |
| DH1325 | 2.335 | 10.000 | 10.000 | 2.695 | 10.000 | 4.717 | 10.000 | 10.000 | 10.000 | 10.000 | 2.994 | 10.000 | 50.000 | 50.000 |
| DH1340 | 10.000 | 10.000 | 10.000 | 10.000 | 10.000 | 10.000 | 10.000 | 10.000 | 10.000 | 10.000 | 10.000 | 10.000 | 26.000 | 50.000 |
| DH1335 | 10.000 | 10.000 | 10.000 | 10.000 | 10.000 | 10.000 | 10.000 | 10.000 | 10.000 | 10.000 | 10.000 | 10.000 | 41.000 | 50.000 |
| DH1338 | 0.020 | 0.425 | 0.307 | 0.061 | 0.040 | 0.067 | 0.067 | 0.062 | 0.064 | 0.063 | 0.061 | 0.060 | 40.000 | 50.000 |
| DH1328 | 0.321 | 10.000 | 1.081 | 2.041 | 0.560 | 1.957 | 3.236 | 3.040 | 1.931 | 2.083 | 1.815 | 2.933 | 25.000 | 50.000 |
| DH1337 | 0.202 | 2.972 | 1.178 | 0.602 | 0.320 | 0.664 | 0.716 | 0.716 | 0.561 | 0.671 | 0.652 | 0.676 | 50.000 | 50.000 |
| DH1324 | 0.160 | 10.000 | 10.000 | 0.613 | 0.650 | 0.880 | 0.917 | 0.908 | 0.620 | 0.759 | 0.788 | 0.879 | 50.000 | 50.000 |
| DH1336 | 0.454 | 10.000 | 10.000 | 0.536 | 0.790 | 0.807 | 1.272 | 1.458 | 0.551 | 0.843 | 0.577 | 1.311 | 50.000 | 50.000 |
| DH1326 | 1.587 | 10.000 | 10.000 | 1.299 | 0.760 | 1.460 | 2.004 | 1.669 | 1.350 | 1.748 | 1.689 | 2.169 | 50.000 | 50.000 |
| DH1339 | 10.000 | 10.000 | 10.000 | 10.000 | 10.000 | 10.000 | 10.000 | 10.000 | 10.000 | 10.000 | 10.000 | 10.000 | 50.000 | 50.000 |
| DH1331 | 10.000 | 10.000 | 10.000 | 10.000 | 10.000 | 10.000 | 10.000 | 10.000 | 10.000 | 10.000 | 10.000 | 10.000 | 50.000 | 50.000 |
| DH1327 | 10.000 | 10.000 | 10.000 | 10.000 | 10.000 | 10.000 | 10.000 | 10.000 | 10.000 | 10.000 | 10.000 | 10.000 | 50.000 | 50.000 |
| DH1332 | 0.601 | 10.000 | 10.000 | 0.674 | 10.000 | 2.370 | 10.000 | 2.809 | 10.000 | 10.000 | 10.000 | 10.000 | 50.000 | 50.000 |
| DH1334 | 10.000 | 10.000 | 10.000 | 10.000 | 10.000 | 10.000 | 10.000 | 10.000 | 10.000 | 10.000 | 10.000 | 10.000 | 50.000 | 50.000 |
| DH1342 | 10.000 | 10.000 | 10.000 | 10.000 | 10.000 | 10.000 | 10.000 | 10.000 | 10.000 | 10.000 | 10.000 | 10.000 | 50.000 | 50.000 |

<0.01 | <0.1 | <1 | <10 | >10

<0.01 | <0.1 | <1 | <10 | <50 | >50

# C

| Group | mAb | BA.4/BA.5 | BA.4.6 | BA.2.75.2 | BF.7 | BQ.1.1 | XBB.1 | XBB1.5 |
|---|---|---|---|---|---|---|---|---|
| Outer RBD face macaque antibodies | DH1333 | 0.030 | 50.000 | 50.000 | 50.000 | 50.000 | 50.000 | 50.000 |
| | DH1345 | 3.400 | 1.700 | 6.300 | 1.500 | 4.400 | 10.000 | 8.500 |
| | DH1341 | 12.000 | 7.100 | 6.500 | 4.300 | 5.700 | 5.300 | 10.000 |
| | DH1344 | 21.000 | 11.000 | 11.000 | 11.000 | 13.000 | 43.000 | 20.000 |
| | DH1343 | 25.000 | 7.100 | 13.000 | 4.900 | 6.100 | 20.000 | 12.000 |
| | DH1329 | 50.000 | 50.000 | 50.000 | 50.000 | 50.000 | 50.000 | 50.000 |
| | DH1340 | 50.000 | 50.000 | 50.000 | 29.000 | 50.000 | 20.000 | 34.000 |
| | DH1335 | 50.000 | 50.000 | 47.000 | 50.000 | 50.000 | 18.000 | 39.000 |
| | DH1330 | 50.000 | 50.000 | 50.000 | 50.000 | 50.000 | 50.000 | 50.000 |
| Inner RBD face macaque antibodies | DH1338 | 50.000 | 50.000 | 50.000 | 50.000 | 50.000 | 50.000 | 50.000 |
| | DH1328 | 50.000 | 45.000 | 50.000 | 35.000 | 50.000 | 39.000 | 50.000 |

$IC_{50}$ (ug/mL): <0.01 | <0.1 | <1 | <10 | <50 | >50

**Figure S8. Distinct antibody clones confer pan-SARS-CoV-2 variants of concern neutralization or Sarbecovirus cross-neutralization.** (A) Vaccine-induced macaque antibody concentration required to block 50% of human antibody binding to Spike. (B,C) IC50 neutralization titer for monoclonal antibodies isolated from wild-type mRNA-LNP vaccinated rhesus macaques. Titers are shown for replicating SARS-CoV-2, SARS-CoV, RsSCHC014 viruses, and SARS-CoV-2 pseudovirus variants of concern. A subset of antibodies were tested against pseudoviruses of SARS-CoV-2 Omicron sublineages (BA.4/BA.5, BA.4.6, BA2.75.2, BF.7, BQ.1.1, XBB.1 AND XBB1.5) variants in 293T-ACE2 cells.

### Supplementary figure 9

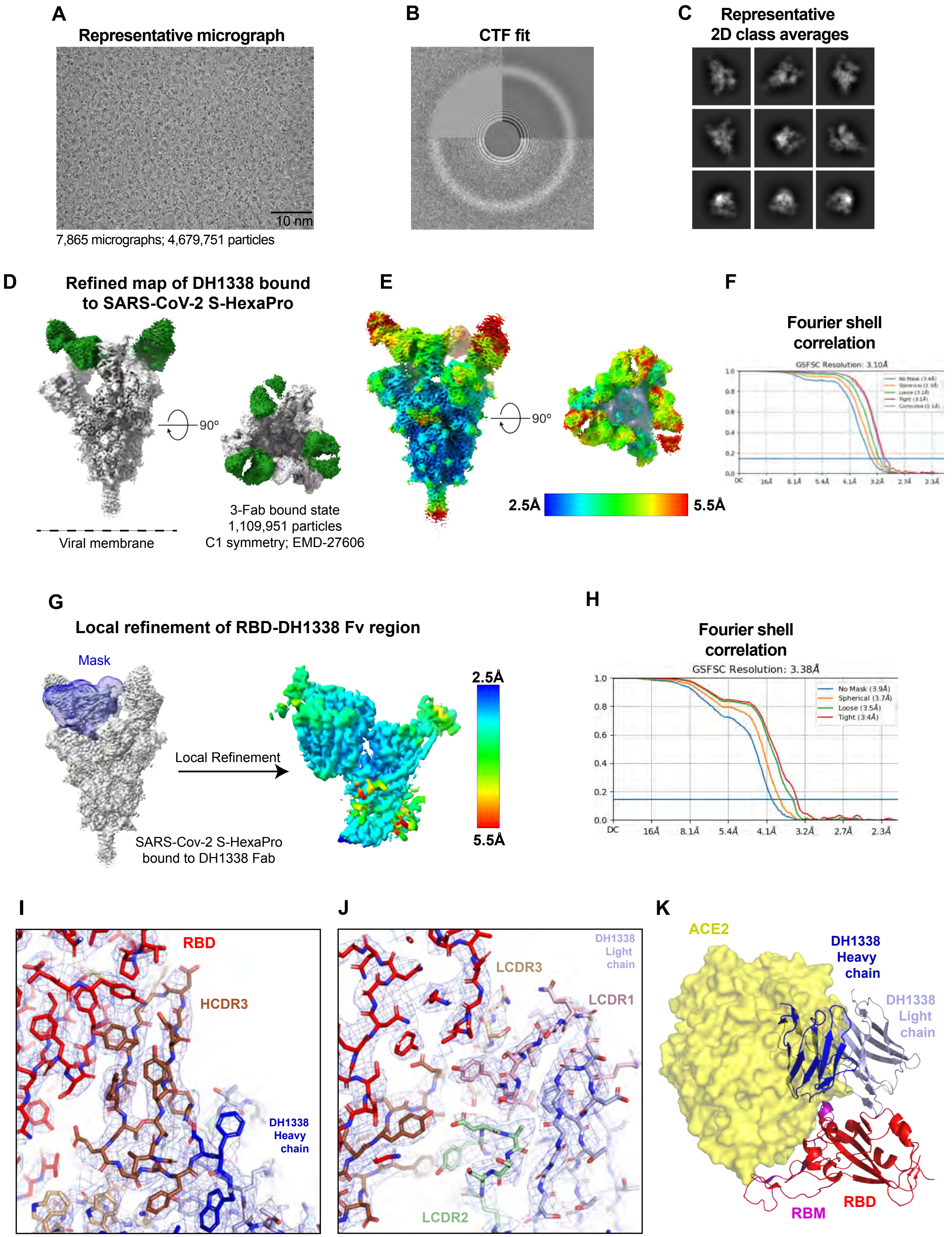

**Figure S9. Cryo-EM data processing for Antibody DH1338 in complex with SARS-CoV-2 S protein, Related to Figure Cryo-EM-1.** (A) Representative micrograph. (B) Representative CTF fit. (C) Representative 2D class averages from cryo-EM dataset. Box size = 324 Å. (D) Refined 3D map segmented and colored by component, with the SARS-CoV-2 S-HexaPro protein colored in grey and DH1338 colored green. (E) Refined map shown in D colored by local resolution. (F) Fourier Shell Correlation (FSC) curves of the 3D reconstruction shown in D with horizontal blue line indicating FSC 0.143. (G) Local refinement of RBD-DH1338 Fv region. Left Blue mesh shows the mask that was used for local refinement. Right. Local Refined 3D density map colored by local resolution. (H) FSC curves of local refined map shown in G. (I) DH1338 HC binding interface with RBD, shown in sticks with electron density shown in blue mesh. (J) DH1338 LC binding interface with RBD. PDBID: 8DPZ. (K) ACE2 (yellow surface representation; PDB 6M0J) binding to RBD (Red cartoon with RBM shown in purple; PDB 6M0J) is sterically hindered by DH1338 (Blue, HC; light blue, LC; cartoon representation; PDB 8DPZ).

### Supplementary figure 10

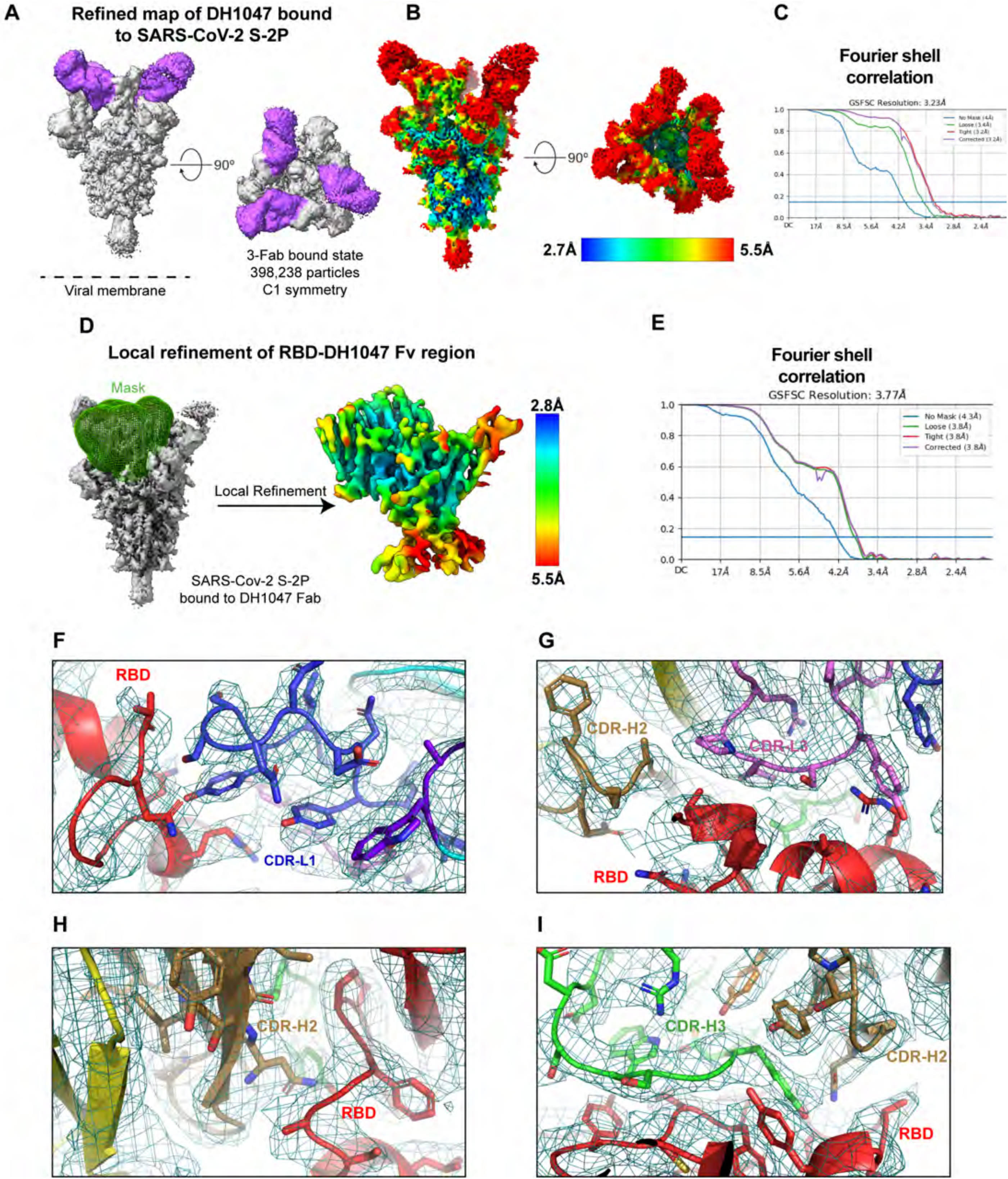

**Figure S10. Cryo-EM data processing for Antibody DH1047 in complex with SARS-CoV-2 S protein.** (A) Refined map of the SARS-CoV-2 2P spike bound to three DH1047 Fabs, highlighted in violet. (B) The map from A. colored by local resolution. (C) Fourier Shell Correlation for the refinement of the full spike-Fab complex. (D) Local refinement region covered by the mask shown in green mesh and resulting map from local refinement, colored by local resolution. The resulting map included the spike RBD and Fv region of DH1047 (E) Fourier Shell Correlation of the locally refined map. (F-**I**) Representative views of the fit of the model to the local refinement map. (F) CDR-L1 to RBD contact. (G) CDR-L3 and H2 to RBD contact. (H) CDR-H2 to RBD contact. (**I**) CDR-H2 and H3 to RBD contact.

**Table S1. Detailed interactions of DH1338 HC with RBD from Pisa Server.**

| | Residue | HSDC | BSA |
|---|---|---|---|
| DH1338 Heavy chain | F:ARG 31 | H | 69.03 |
| | F:SER 53 | | 10.41 |
| | F:GLU 99 | H | 19.12 |
| | F:ASP 100 | | 1.23 |
| | F:ASP 100A | H | 35.04 |
| | F:TYR 100B | H | 99.75 |
| | F:GLY 100C | | 39.88 |
| | F:TYR 100D | H | 121.83 |
| | F:TYR 100E | | 48.20 |
| | F:TYR 100F | H | 31.61 |
| | F:GLU 100H | H | 65.90 |
| RBD | B:TYR 369 | H | 68.21 |
| | B:ASN 370 | | 18.17 |
| | B:SER 371 | H | 16.90 |
| | B:ALA 372 | | 44.44 |
| | B:PHE 374 | | 23.44 |
| | B:SER 375 | H | 59.37 |
| | B:THR 376 | H | 20.93 |
| | B:PHE 377 | H | 41.65 |
| | B:LYS 378 | | 39.03 |
| | B:CYS 379 | H | 29.48 |
| | B:TYR 380 | | 2.98 |
| | B:GLY 381 | | 22.73 |
| | B:VAL 382 | | 5.28 |
| | B:SER 383 | H | 35.93 |
| | B:PRO 384 | | 26.83 |
| | B:THR 385 | | 4.82 |
| | B:ASN 437 | | 3.14 |
| | B:VAL 503 | | 30.77 |
| | B:TYR 508 | H | 13.46 |

HSDC: Residues making Hydrogen/Disulphide bond, Salt bridge or Covalent link, BSA: Buried Surface Area, Å2. |||| Buried area percentage, one bar per 10%.

**Table S2. Detailed interactions of DH1338 LC with RBD from Pisa Server.**

<table>
<tr><th></th><th>Residue</th><th>HSDC</th><th>BSA</th><th></th></tr>
<tr><td rowspan="5">DH1338 Light chain</td><td>G:SER 28</td><td></td><td>11.51</td><td>||</td></tr>
<tr><td>G:SER 30</td><td>H</td><td>57.51</td><td>|||||||||</td></tr>
<tr><td>G:SER 31</td><td></td><td>19.40</td><td>|||||</td></tr>
<tr><td>G:TYR 32</td><td>H</td><td>69.75</td><td>|||||||</td></tr>
<tr><td>G:SER 67</td><td></td><td>0.25</td><td>|</td></tr>
<tr><td rowspan="11">RBD</td><td>B:ARG 403</td><td></td><td>0.58</td><td>|</td></tr>
<tr><td>B:GLY 404</td><td>H</td><td>10.12</td><td>||||||</td></tr>
<tr><td>B:ASP 405</td><td>H</td><td>39.62</td><td>||||||</td></tr>
<tr><td>B:ARG 408</td><td></td><td>8.58</td><td>|</td></tr>
<tr><td>B:THR 500</td><td></td><td>11.05</td><td>|</td></tr>
<tr><td>B:GLY 502</td><td></td><td>32.48</td><td>||||||||</td></tr>
<tr><td>B:VAL 503</td><td>H</td><td>59.62</td><td>|||||||</td></tr>
<tr><td>B:GLY 504</td><td></td><td>26.08</td><td>||||||||||</td></tr>
<tr><td>B:TYR 505</td><td></td><td>25.69</td><td>|||</td></tr>
<tr><td>B:GLN 506</td><td></td><td>8.88</td><td>||||</td></tr>
<tr><td>B:TYR 508</td><td></td><td>1.52</td><td>|</td></tr>
</table>

HSDC: Residues making Hydrogen/Disulphide bond, Salt bridge or Covalent link, BSA: Buried Surface Area, Å2. |||| Buried area percentage, one bar per 10%.

**Table S3. DH1047-Bound SARS-CoV-2 S-2P: Local Refinement of RBD/Fab interface.**

| Cryo-EM data collection and refinement statistics. | |
|---|---|
| **Structure Name** | **DH1047-Bound SARS-CoV-2 S-2P: Local Refinement of RBD/Fab interface** |
| **PDB ID** | 8DTK |
| **EMDB ID** | EMD-27703 |
| **Data Collection and processing** | |
| Microscope | FEI Titan Krios |
| Detector | Gatan K3 |
| Magnification | 81000 |
| Voltage (kV) | 300 |
| Electron exposure (e-/Å^2) | 66.77 |
| Defocus Range (μm) | ~0.75-2.50 |
| Pixel size (Å) | 1.058 |
| Reconstruction software | cryoSPARC |
| Symmetry imposed | C1 |
| Initial particle images (no.) | 2,797,281 |
| Final particle images (no.) | 398,238 |
| Map resolution (Å) | 3.77 |
| FSC threshold | 0.143 |
| **Model composition** | |
| Nonhydrogen atoms | 6942 |
| Protein residues | 449 |
| **R.M.S. deviations** | |
| Bond lengths (Å) | 0.005 |
| Bond angles (°) | 1.133 |
| **Validation** | |
| MolProbity score | 1.53 |
| Clashscore | 4.32 |
| Rotamer Outliers (%) | 0 |
| **Ramachandran plot** | |
| Favored regions (%) | 95.49 |
| Allowed (%) | 4.51 |
| Disallowed regions (%) | 0 |
